# Geometry–Encoded Microtrenches Stabilize Endothelium on High Shear Biomaterial Surfaces

**DOI:** 10.64898/2026.03.16.712222

**Authors:** Aminat M Ibrahim, George Zeng, Scott J Stelick, James F Antaki, Jonathan T Butcher

## Abstract

Maintaining a confluent, antithrombotic endothelium on cardiovascular biomaterial surfaces remains a major barrier to long-term hemocompatibility, as endothelial cells (ECs) rapidly denude under supraphysiological shear in prosthetic devices. Here, we hypothesized that mesoscale surface geometry (∼100–200 µm) could reorganize near-wall hemodynamics, preserving endothelial coverage and function under extreme shear. Engineered microtrenches were introduced onto an implant biomaterial to generate spatially defined shear environments.

Under supraphysiological near-wall shear (∼250 dyn/cm²), microtrenched geometries created attenuated shear and vorticity gradients. Endothelial monolayers were sustained in these flow domains for 120 hours, whereas flat controls rapidly denuded. Endothelial retention in 22.5° angled trenches increased dramatically, from an EC₅₀ of 33 to 101 dyn/cm². 45° angled trenches further increased endothelial shear resistance to an EC₅₀ of 207 dyn/cm².

Endothelial monolayers demonstrated collective mechano-adaptation to ultra-high shear through VE-cadherin junction thickening and coordinated cytoskeletal and nuclear alignment. Mechanoadapted monolayers exhibited increased eNOS expression correlated with local shear and elevated nitrite production (45°: 50.4 ± 6.1 µM; 22.5°: 35.7 ± 3.3 µM; 0°: 28.4 ± 6.8 µM). In contrast, interfaces with abrupt shear transitions or elevated rotational flow exhibited reduced coverage, junctional thinning, and re-emergence of VCAM-1 and PAI-1, indicating inflammatory and pro-thrombotic activation.

Structural, functional, and inflammatory readouts exhibited peak responses within a shared shear–vorticity regime. Multivariate regression identified shear–vorticity coupling as the dominant predictor of endothelial persistence, with optima clustering within a mechanical range (≈0.8–2.9 × 10⁶ dyn·cm⁻²·s⁻¹).

These findings establish geometry-driven modulation of near-wall flow as a predictive, material-agnostic strategy for endothelialization and vasoprotection of high-shear cardiovascular implants.

## Introduction

Despite significant advances in cardiovascular device design, achieving durable hemocompatibility remains one of the field’s most persistent challenges [1,2]. Mechanical heart valves, ventricular assist devices, vascular grafts, and endovascular implants continue to trigger thrombo-inflammatory reactions at the blood-material interface, necessitating lifelong systemic anticoagulation [3–5]. Although anticoagulation mitigates device-induced thrombosis, it exposes patients to significant hemorrhagic risk and limits access to these technologies in low-resource settings where monitoring is restricted [6]. There is a critical clinical need for surface technologies that restore the blood-contacting interface to a native, non-thrombogenic state without the need for systemic pharmacological intervention.

Endothelialization theoretically ensures native antithrombotic and anti-inflammatory surface [7–9]. Yet maintaining a cohesive endothelial monolayer that preserves junctional integrity and vasoprotective signaling under supraphysiological and spatially heterogeneous shear stresses characteristic of mechanical devices, often exceeding 250 dyn/cm², has remained an elusive goal [10–12]. Endothelial cells are highly sensitive to local flow organization. Physiological laminar shear promotes junctional integrity and nitric oxide (NO) signaling, a key mediator of vasoprotective and antithrombotic endothelial function. These phenotypes are typically associated with a physiological shear range of approximately 10–50 dyn/cm² [13,14]. In contrast, the extreme or disturbed flow profiles found in mechanical implants promote junctional fragmentation, inflammatory activation, and eventual cell detachment [15].

Current surface engineering approaches aimed at reducing thrombogenicity have focused predominantly on biochemical and biomolecular surface modifications, such as nitric oxide-releasing polymers that locally deliver NO to inhibit platelet activation [16], heparinized layers [17], peptide-functionalized substrates [18], extracellular matrix coatings [19], and active cell-retention strategies using magnetic nanoparticles [20].

Surface nanotopographies and microscale features below 10 µm have also been explored to enhance endothelial adhesion under flow [21–23]. While these strategies can enhance cell adhesion and improve short-term compatibility [24–27], they do not substantially modify the near-wall shear environment experienced by adherent endothelial monolayers under supraphysiological flow and therefore may not address the flow conditions that drive endothelial detachment.

Endothelial cells integrate both flow-derived and substrate-derived stresses within their microenvironment, including geometry, force distribution, and spatial confinement, to regulate adhesion and phenotype [28,29]. In native vessels, spatially organized and unidirectional flow supports endothelial alignment and junctional organization [29,30]. Consistent with this principle, emerging evidence demonstrates that local surface geometry can actively influence endothelial organization and thrombogenic responses rather than serving as a passive scaffold [31,32]. When steep shear gradients develop, endothelial monolayers exhibit dysfunction and detachment [33]. Collectively, these studies establish the importance of mechanical microenvironmental design in biomaterial surfaces.

Prior microfabricated trench designs from our group demonstrated that geometric modulation of prosthetic surfaces can locally attenuate shear stress under ultra-high flow conditions [34]. However, these early designs were constrained to simple rectangular geometries imposed by fabrication limitations. Although supraphysiological stresses were reduced, extensive regions experienced sub-physiological shear (<10 dyn/cm²). Because endothelial homeostasis requires exposure to a defined shear window (∼10–50 dyn/cm²) [35], these designs provided only limited regions within the vasoprotective range. Subsequent computational fluid dynamics (CFD) analyses predicted that angled trench walls could sustain broader regions of shear, thereby enhancing endothelial coverage and reducing thrombogenic deposition [36]. Building on these advances, we hypothesized that mesoscale microtrenches could organize near-wall hemodynamics into multicellular flow niches capable of preserving endothelial monolayer integrity and nitric oxide signaling under supraphysiologic shear.

Here, we evaluate how mesoscale microtrench geometries regulate local near-wall hemodynamics and determine endothelial structural and functional responses under supraphysiological shear. By integrating CFD with quantitative biological analyses, we identify the hemodynamic conditions that distinguish flow-driven denudation from endothelial persistence. These findings establish a geometry-encoded, coating-free design principle for engineering biomaterial surfaces exposed to supraphysiologic shear.

## Materials and Methods

### Microtrench Fabrication and Profilometry

Microtrench geometries (0°, 22.5°, 45°) were embossed into 1 mm-thick medical-grade UHMWPE sheets using a precision-machined copper mold under controlled pressure and temperature to generate features approximately 80 μm wide and 200 μm deep. Samples were sequentially rinsed in ethanol and PBS prior to cell culture. Surface topography was characterized using a Keyence VK-X260 3D laser-scanning confocal microscope, which provided trench height (hT), width (wT), peak width (wP), and angle (θT). Height maps were exported as STL and CSV files for computational mesh generation (Fig. 1a).

**Fig. 1.**
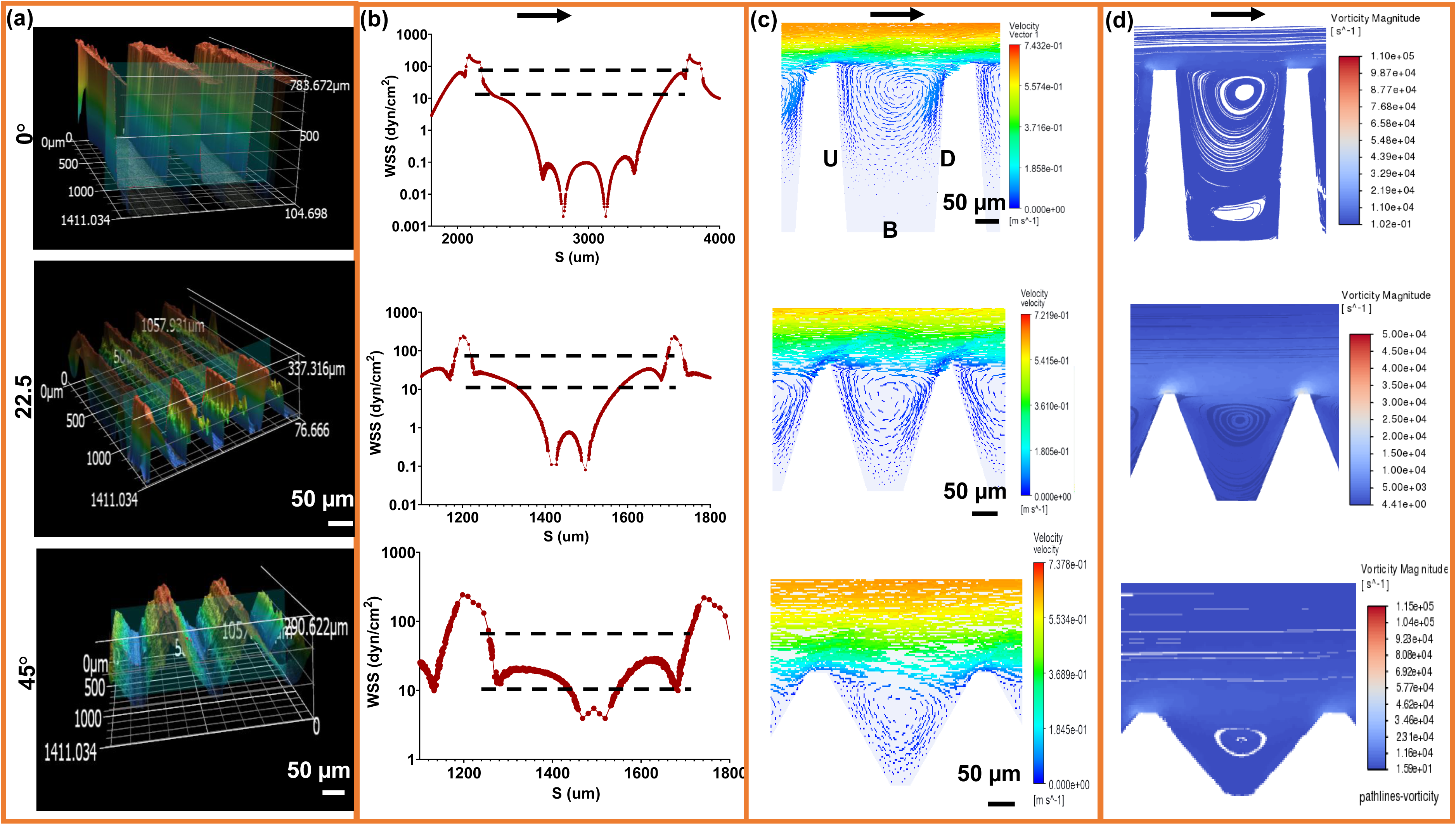
Microtrenches create discrete shear niches and recirculation zones. (a) 3D profilometry and computational reconstructions of 0°, 22.5°, and 45° trench geometries designed to impose controlled flow gradients. (b) Simulated wall shear stress (WSS) distributions reveal geometry-dependent heterogeneity, with high-shear ridges and ultra-low-shear regions at trench bases. Dashed lines indicate the physiological WSS range that supports endothelial quiescence. (c) Velocity vector contours illustrate flow acceleration along the trench peaks and localized separation along the slopes, generating distinct shear microenvironments. (d) Vorticity magnitude maps highlight rotational cores and recirculating eddies concentrated at the trench bases, marking potential loci for disturbed flow and mechanotransductive signaling. Arrows denote flow direction. Scale bars, 50 µm.

### Computational Fluid Dynamics

CFD models were generated in ANSYS Fluent using profilometry-derived trench cross-sections extruded into 3D to a constant spanwise thickness, forming a three-trench segment that captures near-wall flow organization while remaining quasi-2D in the spanwise direction. Media was modeled as a Newtonian fluid (density 1000 kg·m⁻³, viscosity 0.00095 Pa·s). At the imposed shear rates and laminar flow regime considered here, viscosity variations are minimal, enabling consistent comparison across trench geometries. A uniform inlet velocity was prescribed to achieve a near-wall bulk shear of approximately 250 dyn/cm², matching bioreactor conditions. Laminar, steady-state flow with no-slip boundary conditions was assumed.

Simulations were performed using a pressure-based steady-state solver with second-order spatial discretization. Convergence was confirmed at residuals below 10⁻⁶, and wall shear stress values stabilized. Mesh refinement near trench walls resolved gradients in WSS, vorticity, WSSG, and flow direction, and mesh independence was verified (Supplementary Table 4). Reynolds numbers remained within the laminar regime under all conditions.

CFD-derived upstream, base, and downstream regions were mapped onto confocal images to define biologically analyzed zones.

### Endothelial Cell Isolation and Culture

Porcine aortic valvular endothelial cells (PAVECs) were isolated by collagenase digestion as described previously [37]. Cells were expanded in Dulbecco’s Modified Eagle’s Medium (DMEM, Fisher Scientific) supplemented with 10% fetal bovine serum (GeminiBio) and 1% penicillin-streptomycin (Gibco). Cells between passages 4 and 5 were used. Before seeding, UHMWPE substrates were coated with 50 μg/ml collagen (ENZO) for 1h at 37°C and 5% CO2. Cells were seeded at 100,000 cells/cm² onto flat or microtrenched samples and allowed to adhere for 24 hours before flow exposure.

### Flow Bioreactor and Shear Exposure

A custom parallel-plate bioreactor was constructed using a glass flow plate with drilled inlet and outlet ports. UHMWPE samples were seated within a PDMS gasket formed inside a silicone housing, with a precision spacer defining the channel height and flow field. The chamber was connected via tubing to a compliance dampener and driven by a peristaltic pump (Masterflex) at a constant flow rate corresponding to a calculated near-wall shear stress of ∼250 dyn/cm², based on channel height and volumetric flow rate.

Cells were exposed to continuous shear for 48h for primary assays, and for 2 or 5 days for long-term studies to allow for proper PAVEC phenotype expression. All experiments were conducted at 37°C and 5% CO₂.

### Immunofluorescence Staining and Imaging

Samples were fixed in 4% PFA for 20 minutes, permeabilized in 0.2% Triton Solution, and blocked in 3% bovine serum albumin for 1 hour. Primary stains included the following: VE-cadherin (junctional structure; Bio-Rad), eNOS (functional signaling; Cell Signaling), Phalloidin (actin cytoskeleton; Invitrogen), PAI-1 and VCAM-1 (activation markers; Invitrogen), and DAPI (nuclei; Thermo Fisher), incubated overnight at 4 °C. Afterwards, secondary antibodies (Alexa Fluor) were incubated for 30 minutes at room temperature. Images were acquired on a Zeiss LSM 880 multiphoton confocal microscope with identical laser and detector settings for all samples.

### Image Processing and Quantification

All image analysis was performed in Fiji (ImageJ v1.54p). CFD-derived upstream, base, and downstream ROIs were applied uniformly across all images.

Endothelial coverage: Phalloidin and VE-cadherin images were converted to 8-bit, contrast-normalized, background-subtracted, and combined. Otsu thresholding generated binary masks. Minor morphological operations were applied only when necessary to maintain mask continuity (e.g., remove objects smaller than 2 μm). CFD-defined ROIs were then overlaid to compute area fraction within each region. (Validation of mask accuracy is reported in Supplementary Methods.)

VE-Cadherin junction thickness: Junction thickness was quantified by drawing perpendicular line profiles across intact borders between adjacent cells within each ROI [38]. Three to five junctions were measured per region using ImageJ’s line profile tool.

Cell shape metrics: Whole-cell outlines were traced manually from VE-cadherin images. An optional Gaussian blur (sigma = 1) was applied to smooth minor corrugations. For each region, three well-spread monolayer cells were traced. Area, perimeter, aspect ratio, circularity, Feret’s diameter, and centroid were recorded using Fit Ellipse and Shape Descriptors. ROIs were saved for reproducibility.

Nuclear polarity: Cell and nuclear centroids were obtained from VE-cadherin and DAPI masks[39]. Polarity was calculated as normalized centroid displacement along the flow axis. Negative polarity denotes upstream displacement.

Actin orientation and coherency: OrientationJ was applied to background-subtracted, lightly blurred actin images. The plugin returned dominant fiber orientation (0-90° relative to horizontal) and coherency (0-1). Directionality histograms (90 bins) were generated as a secondary check.

eNOS, VCAM-1, PAI-1 quantification: Channels were thresholded based on cytoplasmic signal distribution and converted to binary masks. Percent positive area was calculated within CFD-defined ROIs. eNOS signal was normalized to endothelial coverage by dividing eNOS-positive area by EC-covered area.

### Griess Quantification Assay

Conditioned media were collected at the end of each flow experiment and analyzed using the Griess Reagent Kit (Invitrogen). Absorbance at 548 nm was measured on a microplate reader, and nitrite concentration was calculated from sodium nitrite standards.

### Statistical Analysis

Statistical analyses were performed in GraphPad Prism (v10.6.1). Normality was assessed using Shapiro-Wilk testing prior to parametric analysis. Significance was set at α = 0.05. One-way ANOVA with Tukey post-hoc testing was used for region-wise comparisons within geometries, and two-way ANOVA was used to evaluate the effects of geometry and region or condition, where applicable. Endothelial coverage values were logit-transformed prior to regression analyses. Multivariate regression models examined the contributions of wall shear stress (WSS), vorticity, wall shear stress gradient (WSSG), and flow direction. Receiver operating characteristic (ROC) analysis was used to evaluate the ability of shear-derived metrics to classify endothelial retention ≥50%. Full model equations and derivations are provided in Supplementary Methods. Data are presented as mean ± SEM unless otherwise stated.

## Results

### Engineered Microtrenches Create Spatially Distinct Shear Niches and Recirculation Zones

To determine how engineered surface geometry modulates local hemodynamics, we fabricated microtrenched substrates with trench angles of 0±4°, 22.5±3°, and 45±3°, informed by computational studies of trenched flows, and designed to generate geometry-defined near-wall shear niches, consistent with prior analyses [36]. Three-dimensional profilometry verified dimensional fidelity and repeatability, providing measured contours for CFD simulations (Fig. 1a). Notably, angled trench geometries are not readily produced using conventional photolithography or etching approaches; their realization here was enabled through precision embossing of UHMWPE.

CFD analysis showed that all trenches locally attenuated wall shear stress (WSS) relative to the applied near-wall shear of ≈250 dyn/cm², reducing intra-trench WSS by more than an order of magnitude. Across the surfaces, WSS ranged from >200 dyn/cm² at ridge crests to ≈ 0.01–5 dyn/cm² within basal pockets, creating a broad shear distribution that includes regions approaching physiological shear levels (∼10–50 dyn/cm²) and forming discrete shear niches separated by gradients spanning two orders of magnitude (Fig. 1b-d), which create transitional flow environments between low- and high-shear regions. Increasing trench angle compressed low-shear regions and amplified ridge-scale acceleration.

Flow organization differed across geometries. The 0° trenches exhibited vertically stacked vortices spanning most of the trench depth, producing near-stagnant recirculation. The 22.5° design generated alternating acceleration-deceleration bands with a single asymmetric downstream vortex, whereas the 45° trenches produced smaller, more confined recirculation zones with sharper forward-flow acceleration, consistent with earlier flow reattachment along steeper walls and the absence of secondary vertical eddies.

Vorticity maps showed rotational shear concentrated at reattachment zones and downstream walls, while forward-flow regions exhibited low vorticity. Pressure mapping confirmed static pressure buildup within recirculation pockets (Supplementary Fig. 1a). Using literature-based hemodynamic classification validated in prior endothelial studies (Supplementary Fig. 1b), surface regions were categorized as low (<10 dyn/cm²), intermediate (10–43 dyn/cm²), or high (>43 dyn/cm²) shear.

Increasing trench angle progressively redistributed surface area from ultra-low shear recirculation toward intermediate and front-facing shear regions. Consistent with computational design predictions, angled trench geometries increased the fraction of surface area within the physiological shear window relative to rectangular trenches. Quantitative surface-area analysis confirmed this redistribution: the fraction of ultra-low shear regions decreased from 75.0% in rectangular trenches to 48.2% and 19.6% in the 22.5° and 45° geometries, respectively, while the fraction of intermediate-shear regions increased from 14.3% to 46.3% and 64.0%. Intra-trench flow fields were spatially smooth and reproducible within each geometry, with dominant differences arising between trench angles.

These results demonstrate that microtrenches generate reproducible continuum of shear and vorticity gradients along surfaces otherwise exposed to high near-wall shear, establishing geometry-defined hemodynamic niches for subsequent endothelial analyses (Fig. 2).

**Fig. 2.**
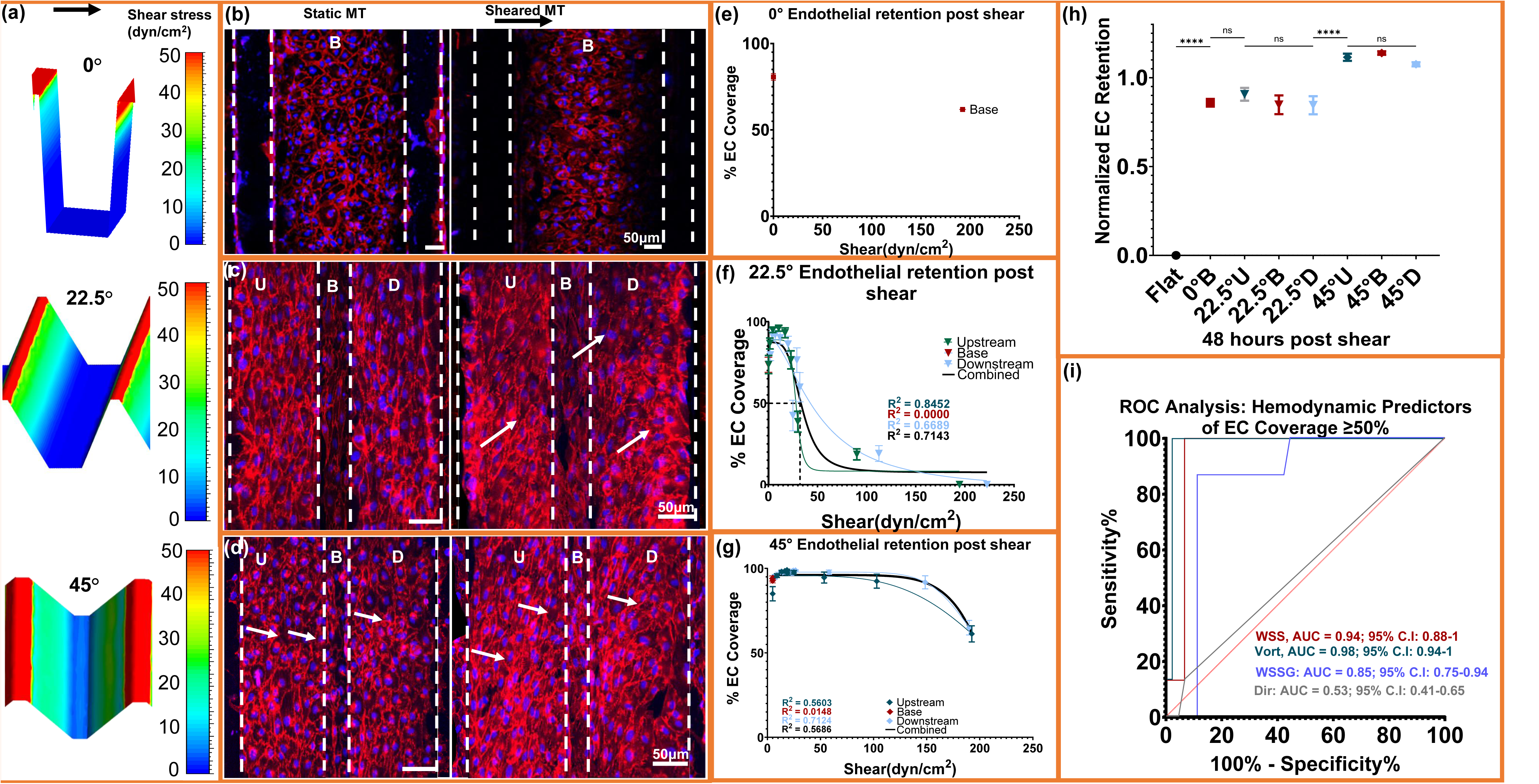
Geometry-defined shear fields sustain endothelial coverage under extreme flow. (a-d), CFD-derived wall shear stress (WSS) distribution map (< 50 dyn/cm²) for 0°, 22.5°, & 45° microtrenches (MTs) and corresponding Immunofluorescence images showing endothelial monolayers under static and 48h post-shear conditions, (VE-cadherin, red; nuclei, blue). (e-g), Quantification of absolute endothelial coverage (% EC projected area fraction) across local shear domains for each geometry with logistic fits (R² shown); Coverage declines monotonically with increasing WSS and remains most preserved in 45°. (h), Geometry-level normalized endothelial retention at 48h (shear/static), computed as post-shear coverage divided by paired static coverage for each region. Significant differences were observed between geometries, with no differences among U, B, and D regions within each geometry. (i) ROC analysis identifying hemodynamic predictors of high EC coverage (≥50 percent coverage). Vorticity provides the strongest discrimination (AUC = 0.98), followed by WSS (AUC = 0.94), WSSG (AUC = 0.85), and flow direction (AUC = 0.53). Data represent mean ± SEM; n = 8 biological samples, with duplicate/triplicate measurements per region per sample. U, B, and D correspond to the Upstream, Base, and Downstream. Coverage values represent projected area fraction within CFD-defined ROIs.

### Flow-patterned microtrenches expand endothelial shear resistance and sustain monolayer retention under supraphysiologic shear

Having established geometry-defined mechanical niches, we next examined how these microenvironments regulate endothelial coverage under sustained flow. Consistent with our prior report [36], endothelial monolayers remained cohesive within microtrenches after shear exposure, whereas flat controls exhibited extensive denudation (Fig. 2b-d).

Geometry-level normalized endothelial retention at 48 h, defined as shear-exposed coverage divided by static coverage, differed significantly between geometries (Flat < 0° ≈ 22.5° < 45°; Fig. 2h), with the 45° design exhibiting the highest retention under supraphysiologic shear. Notably, normalized retention in the 45° geometry exceeded static baseline values (retention >1.0), indicating shear-dependent monolayer expansion rather than simple preservation of initial attachment.

To relate endothelial coverage to local hemodynamics, each geometry was subdivided into upstream, base, and downstream regions using mean WSS (Fig. 2a; Supplementary Fig. 2a-b). Transverse resolution limits precluded resolving cell morphology on the vertical walls of the 0° trench; thus, analysis focused on the flat trench bottom region. The 22.5° and 45° designs spanned low, transitional, and high shear domains, including ridge-peak zones where shear exceeded 200 dyn/cm².

The base of the 0° rectangular trench retained 80.7±2.0% coverage (Fig. 2e) under near-stagnant flow (mean WSS ≈ 0.02 dyn/cm²), serving as a low-shear reference. Across geometries, subregional coverage exhibited a sigmoidal decline with increasing WSS (Fig. 2f, g), and cells remained adherent even at WSS values above 100 dyn/cm². In the 22.5° trenches, upstream and downstream regions followed similar logistic profiles (R² ≈0.67-0.85): coverage remained >70% across low-mid shear (≤15 dyn/cm²) but dropped below 50% in the downstream reversal zone, where steep gradients and recirculation developed. Even forward-flow regions near ∼30 dyn/cm² showed reduced coverage, indicating that elevated vorticity and abrupt shear transitions are associated with reduced endothelial persistence.

The 45° geometry maintained uniformly high coverage and exhibited a shallower sigmoidal decline (combined R² ≈0.57), with nonlinear regression indicating a right-shifted detachment response and a fitted EC₅₀ of approximately 207 dyn/cm². Coverage remained >60% even at ∼190 dyn/cm² near ridge peaks and reached 90-98% across intermediate-shear forward-flow bands (10-43 dyn/cm²). The basal recirculation niche (≈4-6 dyn/cm²) preserved intact monolayers, and transition zones showed only modest losses. Coverage also remained ∼98% in the mild reversal pocket, indicating that smoother gradients and more spatially confined recirculation zones in the 45° design mitigate detachment despite extreme applied shear.

Single-parameter ROC analysis (Fig. 2i) identified vorticity as the strongest discriminator of high coverage regions (AUC = 0.98, p < 0.0001), followed by WSS (AUC = 0.94, p < 0.0001). WSSG showed moderate predictive ability (AUC = 0.85), whereas flow direction performed poorly (AUC = 0.53). These results indicate that local rotational flow structure, particularly when coupled to shear magnitude, is the strongest predictor of endothelial detachment, while gradient steepness and directionality add limited predictive value once rotational intensity is accounted for.

Collectively, these data demonstrate that microtrench geometry organizes discrete hemodynamic niches that define the limits of endothelial persistence, with coupled vorticity and WSS emerging as the dominant predictors of coverage. Regions of reduced coverage localized to high-vorticity or high-gradient interfaces, whereas forward-flow domains sustained coverage even under supraphysiologic shear. These findings establish shear organization and rotational intensity as dominant mechanical determinants of endothelial persistence on engineered surfaces.

### Microtrench shear fields drive collective endothelial monolayer mechano-adaptation

We next examined whether the local shear conditions associated with endothelial coverage also drive coordinated intracellular organization. Cell shape and nuclear polarity exhibited consistent shear-dependent trends. Aspect ratio increased with WSS and plateaued beyond ≈50 dyn/cm², consistent with elongation under sustained forward-flow (Supplementary Fig. 3g). Nuclear polarity indices were predominantly negative in both 22.5° and 45° geometries, indicating upstream nuclear displacement relative to the cell centroid. This upstream bias was strongest in forward-flow regions, whereas reversal and near-stagnant zones showed greater variability (Supplementary Fig. 4a-c). These data indicate that near-wall shear organization, particularly directional coherence, influences endothelial cell elongation and subcellular spatial positioning.

To determine whether these shear-associated morphological patterns corresponded to changes in cell–cell junction structure, we quantified VE-cadherin junction thickness across shear domains (Fig. 3g-i), a known mechano-adaptive response of endothelial junctions to shear stress [15]. Junction thickness exhibited geometry-specific profiles that mirrored the shear niches. In the 22.5° trenches, junctions were thinnest at ≤0.1 dyn/cm² and thickened across the transitional range (≈0.3-10 dyn/cm²), plateauing once shear exceeded physiological levels. In the 45° trenches, junction thickness followed a biphasic profile: thinning under low recirculation shear (≈4-6 dyn/cm²), reinforcement across a broad intermediate shear regime (≈7-103 dyn/cm²), and renewed thinning at extremely high-shear ridge peaks (≈190 dyn/cm²). This secondary thinning at the highest shear levels occurred despite maintained coverage, indicating that junction thickness decreases under extreme shear even when surface coverage is preserved. Upstream and downstream edges behaved similarly within matched shear bins (Supplementary Fig. 3h, i), indicating that junction thickness is associated with local shear magnitude and vorticity structure. Notably, the shear levels that maximized junction thickness coincide with the retention plateau in Fig. 2, linking intermediate shear domains to both increased junction thickness and preserved endothelial coverage. This relationship suggests that junction reinforcement reflects the shear tolerance of the collective endothelial monolayer across the flow niches established by the trench geometries.

**Fig. 3.**
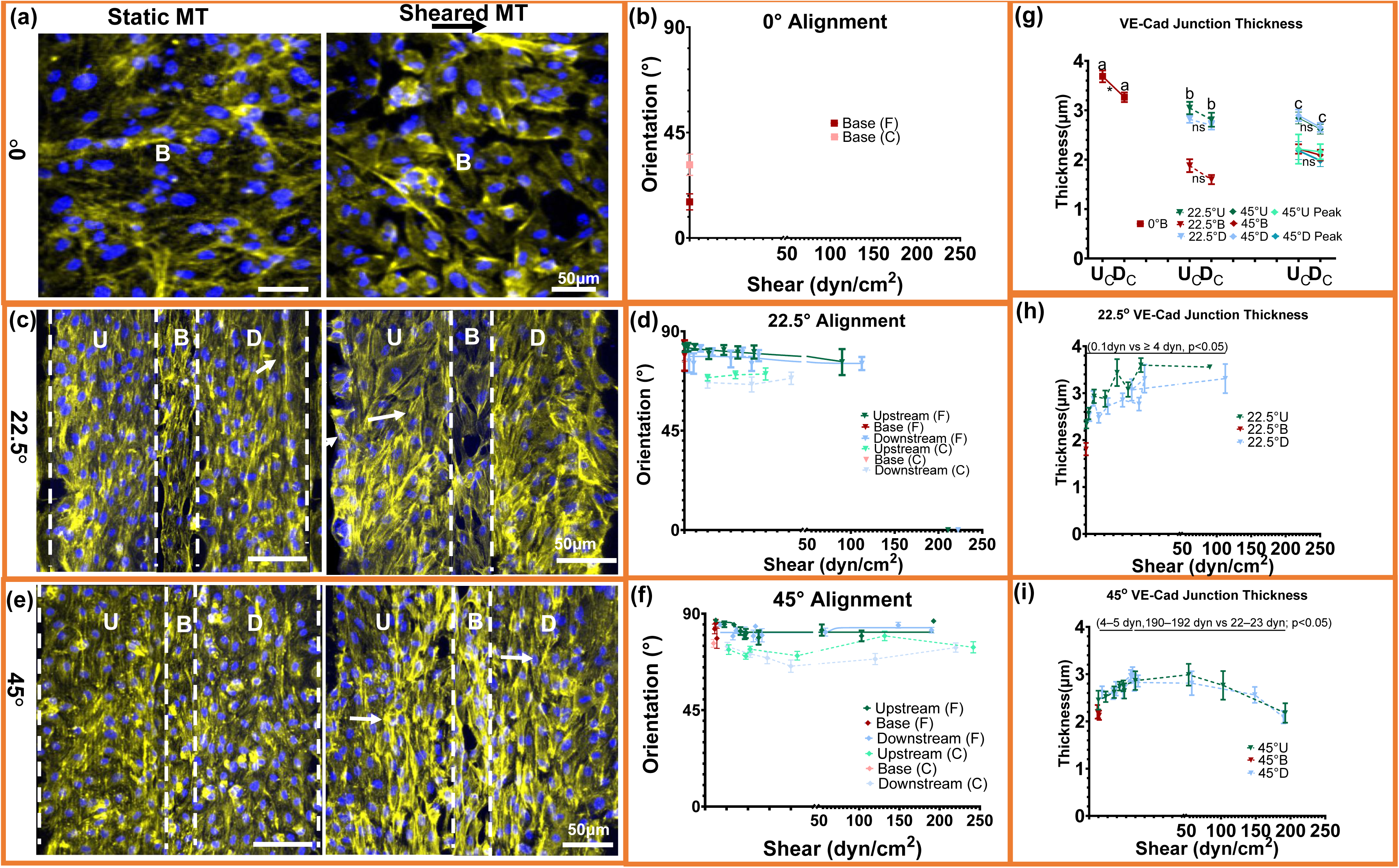
Microtrenches drive coordinated cytoskeletal alignment and VE-cadherin junction remodeling across shear niches. (a, c, e) Immunofluorescence images of endothelial cells within 0°, 22.5°, and 45° microtrenches showing organized actin filaments (yellow) and nuclei (blue) under static and 48 h post-shear conditions. Arrows indicate aligned, elongated morphology. U, B, and D denote Upstream, Base, and Downstream regions. (b, d, f) Quantification of actin-fiber and whole-cell orientation across shear domains. Alignment increased toward perpendicular orientation under forward shear and plateaued beyond ≈50 dyn/cm², with minor curvature-dependent differences between 22.5° and 45° geometries. (g–i) VE-cadherin junction thickness across matched shear domains. Junctions were thinnest in near-stagnant or extreme-shear regions and maximally reinforced within mid-shear, low-vorticity zones (≈22–23 dyn/cm²), corresponding to the retention plateau in Fig. 2. Upstream and downstream edges showed similar responses, indicating dependence on shear magnitude and flow organization rather than flow direction. Data are mean ± SEM; n = 8 biological samples with duplicate/triplicate measurements per region per sample. One-way ANOVA with Tukey’s HSD; **** p < 0.0001, *** p < 0.0005, ** p < 0.005, * p < 0.05, ns = not significant.

Actin filament orientation exhibited similar shear-dependent behavior. Under static conditions, cells displayed disorganized stress fibers, whereas shear exposure induced robust reorientation of both actin filaments and whole-cell major axes nearly perpendicular to flow (≈85-90°) across all geometries (Fig. 3a,c,e). Quantitative analysis showed that mean alignment angles decreased modestly with increasing WSS and plateaued beyond ≈50 dyn/cm² (Fig. 3b,d,f). Actin coherency was lowest in the 0° base (C.I. <0.2) under near-stagnant flow, increased to ≈0.25-0.4 across the 22.5° and 45° trenches, and showed local reductions specifically in high-vorticity or recirculation regions (Supplementary Fig. 3). These regions correspond to areas of reduced coverage in Fig. 2, indicating that decreased cytoskeletal alignment is associated with elevated vorticity and recirculation.

Together, these data show that specific shear domains are associated with upstream nuclear displacement, increased junction thickness within intermediate shear ranges, and perpendicular actin alignment. These structural features co-localize with regions of preserved endothelial coverage under elevated shear, which is associated with shear organization and vorticity structure rather than shear magnitude alone.

### Microtrenched flow niches enhance endothelial eNOS expression and nitric oxide production

Immunofluorescence imaging revealed strong eNOS localization along cell borders and aligned actin filaments after shear (Fig. 4a,c,e), consistent with enhanced nitric oxide signaling. eNOS intensity was highest within flow niches, including the 22.5° mid-slope (≈12-40 dyn/cm²) and the 45° basal region (≈4-6 dyn/cm²), rather than increasing monotonically with shear magnitude. These same areas showed maximal endothelial coverage in Fig. 2. The near-stagnant 0° base displayed diffuse or absent eNOS, consistent with reduced junctional organization and low endothelial coverage.

**Fig. 4.**
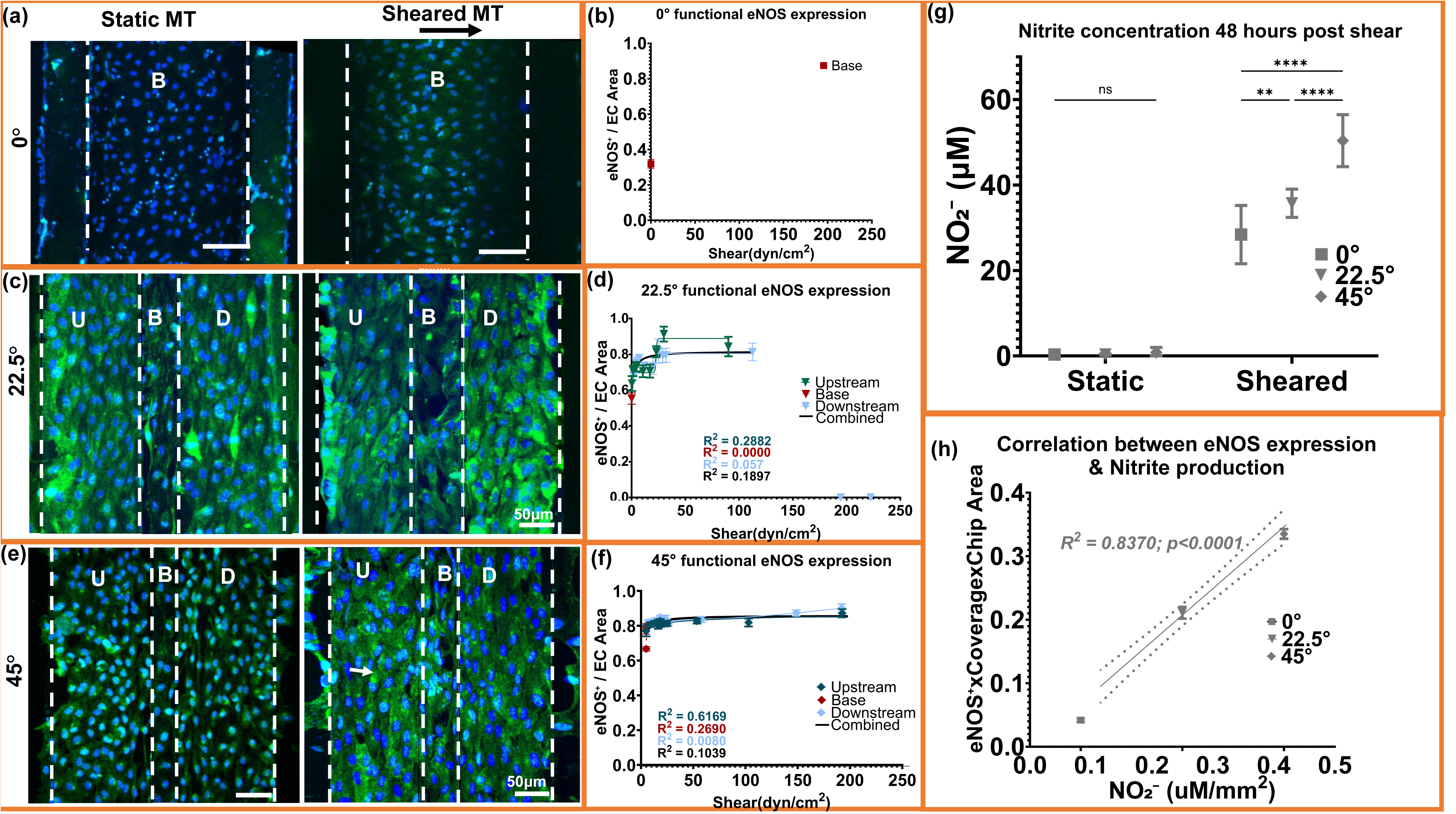
eNOS activation and nitric-oxide production vary across shear niches. (a, c, e) Immunofluorescence images showing endothelial nitric oxide synthase (eNOS, green) and nuclei (blue) in 0°, 22.5°, and 45° microtrenches under static and shear conditions. U, B, and D denote upstream, base, and downstream regions. (b, d, f) Quantified eNOS intensity normalized to endothelial area increased with shear, following a biphasic response across geometries. (g) Nitrite concentration measured 48 h post-shear confirmed elevated nitric-oxide release in all microtrenches compared to static controls, with the 45° geometry showing the highest production. (h) eNOS expression strongly correlated with nitrite output (R² = 0.84, p < 0.0001), confirming direct translation from enzymatic activation to functional release. (i) Quadratic model fitting revealed a parabolic dependence between eNOS and the composite term (WSS × vorticity), with an optimal regime at approximately 1.5 × 10⁶ dyn·cm⁻²·s⁻¹ corresponding to maximal endothelial functionality. Data represent mean ± SEM; n = 8 biological replicates; duplicate/triplicate measurements per region per sample. For nitrite (g): n = 8 (0°), 10 (22.5°), 5 (45°), and measured in SD. One-way ANOVA with Tukey’s HSD test; ****p < 0.0001, ns = not significant.

Quantification normalized to the endothelial area (eNOS %area divided by endothelial coverage within each ROI) confirmed a significant geometry-dependent increase in eNOS (45° > 22.5° > 0°; Fig. 4b,d,f). The response was biphasic, rising sharply across low-mid shear and remaining elevated beyond ≈50-60 dyn/cm². In the 22.5° trenches, eNOS normalized to the endothelial area peaked at ≈20-30 dyn/cm², whereas the 45° geometry sustained high normalized expression across a broader shear range (≈4-60 dyn/cm²). Although the 22.5° trenches produced higher per-area eNOS intensity, the substantially greater monolayer coverage in the 45° design yielded the highest total functional output. The shear levels supporting maximal normalized eNOS expression align with the retention plateau in Fig. 2 and co-localize with shear domains associated with lower vorticity and reduced shear gradients. Regions with elevated vorticity or steep shear transitions exhibited reduced normalized eNOS expression despite increasing shear magnitude.

Griess assay showed increased NO₂⁻ release under shear versus static controls (Fig. 4g), with 45° trenches producing the highest levels (45°: 50.4 ± 6.1 µM; 22.5°: 35.7 ± 3.3 µM; 0°: 28.4 ± 6.8 µM). eNOS intensity strongly correlated with NO₂⁻ concentration (R² ≈ 0.84; Fig. 4h), indicating direct translation from enzymatic activation to metabolic output. In contrast, geometries characterized by recirculation and elevated vorticity exhibited reduced eNOS expression and lower NO₂⁻ release.

Together, elevated eNOS expression and NO₂⁻ production co-localize with shear domains characterized by organized vorticity and preserved endothelial coverage.

### Shear transitioninterfaces and low-shear regions promote pro-inflammatory and pro-thrombotic signaling

Because eNOS expression and NO₂⁻ release were highest within the shear domains identified in Fig. 4, we next asked whether these same mechanical environments also suppress thrombogenic and inflammatory activation. Under static conditions, VCAM-1 and PAI-1 were strongly expressed, most prominently in the near-stagnant base of the 0° trench, but shear exposure reduced their expression across all geometries (Fig. 5a-f). Quantification showed VCAM-1⁺ area falling to <5% and PAI-1⁺ area to <3% in forward-flow regions (Fig. 5g,h), with the strongest suppression occurring in the 22.5° mid-slope and 45° base, precisely the domains that exhibited maximal retention and the highest eNOS activity.

**Fig. 5.**
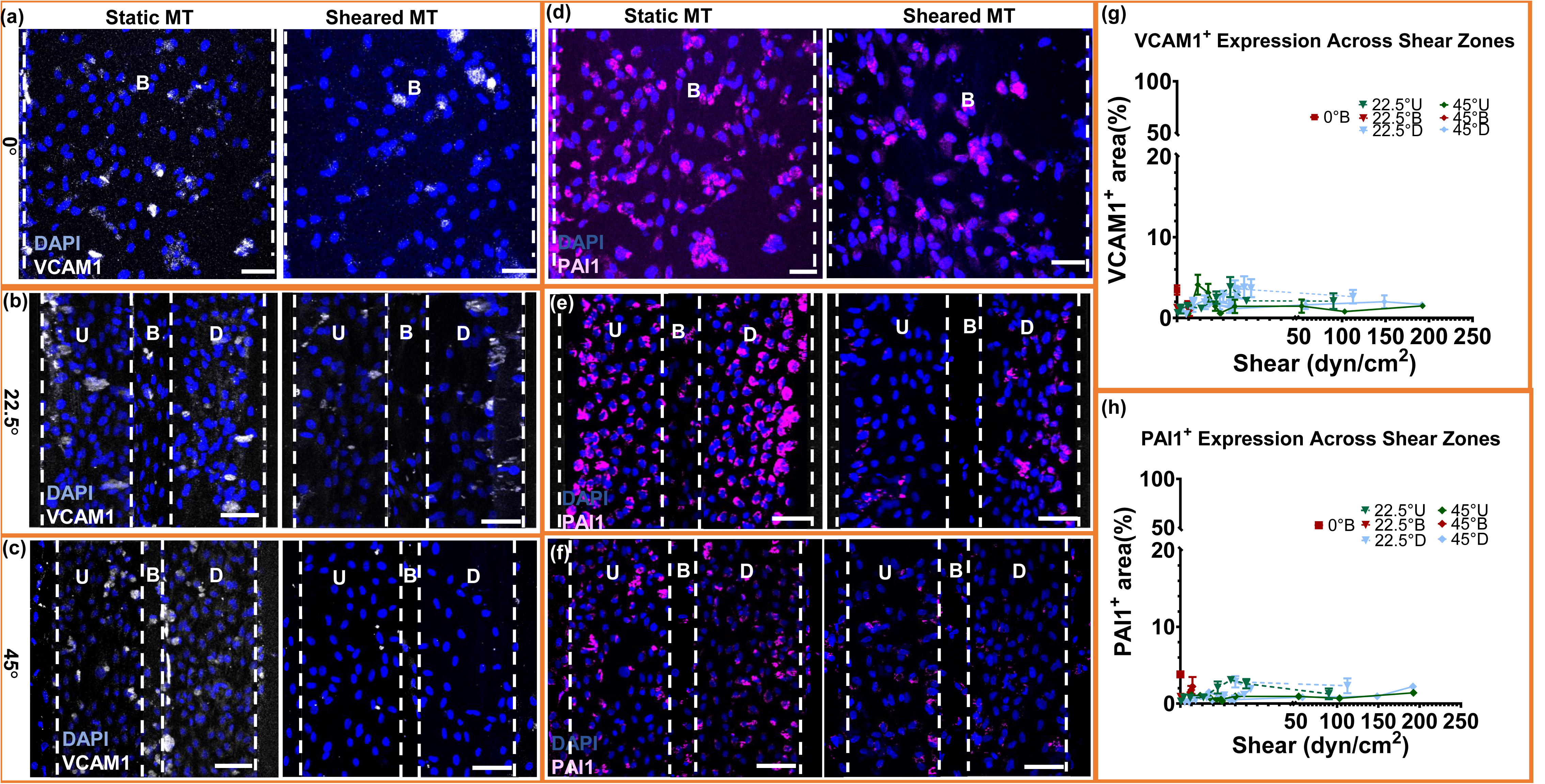
Shear organization suppresses endothelial inflammatory and pro-thrombotic signaling. (a–f) Immunofluorescence staining of ECs cultured on microtrenches (MTs) at 0°, 22.5°, and 45° geometries under static or shear conditions. VCAM-1 (white) and PAI-1 (magenta) were co-stained with DAPI (blue). Dashed lines indicate upstream (U), base (B), and downstream (D) regions within each trench. Scale bars, 50 µm. (g–h) Quantification of VCAM-1⁺ and PAI-1⁺ area (%) across local shear zones, plotted against wall shear stress (dyn/cm²). Data represent mean ± SEM; n = 5 biological replicates per region per sample, with duplicate/triplicate measurements.

Residual VCAM-1 and PAI-1 expression persisted only at flow transition interfaces, including the downstream recirculation zone of the 22.5° trenches and the ridge-peak acceleration region of the 45° geometry. These sites experience strong vorticity or abrupt shear transitions and overlap with regions of reduced eNOS and thinner junctions, indicating that regions with elevated vorticity or steep shear gradients are associated with pro-inflammatory and pro-thrombotic marker expression.

Across all geometries, VCAM-1 and PAI-1 displayed inverse trends relative to eNOS (Figs. 4-5). VCAM-1 was minimized between ∼30-50 dyn/cm² under forward shear, whereas PAI-1 remained suppressed across a broader shear range (∼10-60 dyn/cm²), suggesting differential sensitivity to shear–vorticity organization. Importantly, the regions with the lowest VCAM-1/PAI-1 were the same domains that sustained high retention and maximal eNOS. These results demonstrate that endothelial shear niches supporting elevated eNOS activity are associated with reduced VCAM-1 and PAI-1 expression, whereas high-vorticity shear transition interfaces shift the balance toward pro-inflammatory signaling.

### Endothelial retention and vasoprotective signaling sustained in chronic high shear

To determine whether the shear-dependent responses observed at 48h represent transient remodeling or a stable phenotype, 22.5° microtrenches were exposed to continuous flow for 2 or 5 days (Fig. 6a-f). VE-cadherin staining showed cohesive monolayers at both time points, with day 5 samples displaying fewer discontinuities and fully preserved upstream, base, and downstream regions (Fig. 6a). Actin imaging revealed that the perpendicular fiber alignment established at 48h persisted through day 5 (Fig. 6b). eNOS staining remained robust across all regions (Fig. 6c), consistent with sustained nitric oxide signaling.

**Fig. 6.**
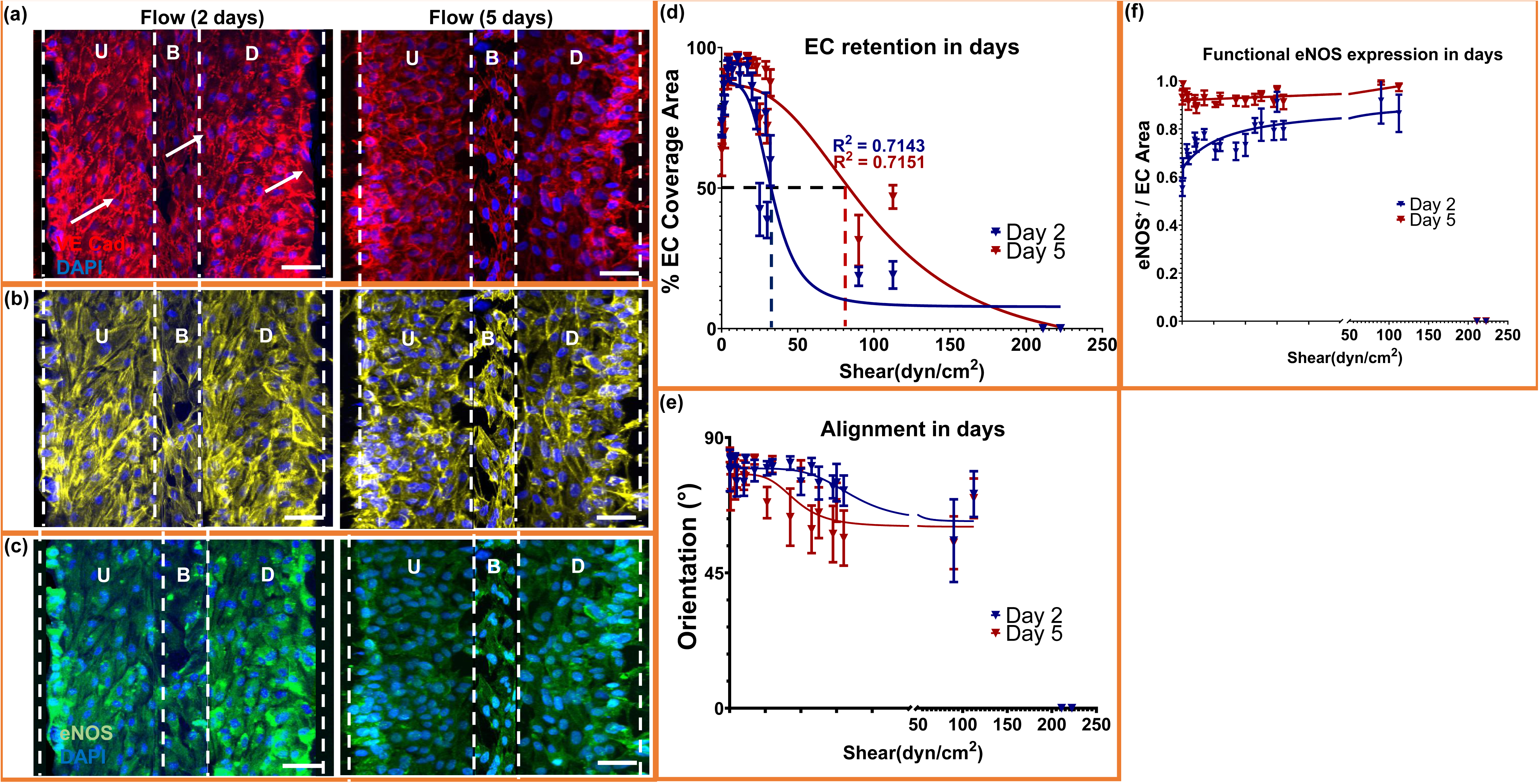
Microtrenches sustain endothelial coverage and mechanoadaptation under chronic shear. (a–c) Immunofluorescence images of 22.5° microtrenches showing endothelial cells cultured under continuous shear for 2 and 5 days. VE-cadherin (red) and DAPI (blue) mark junctional integrity (a), ACTIN denotes alignment (b), and eNOS (green) indicates functional activation (c). U, B, and D correspond to upstream, base, and downstream regions, respectively. (d) Quantification of endothelial coverage versus shear showing sustained retention and a rightward shift in the sigmoidal response after 5 days, indicating increased shear tolerance. (e) Orientation analysis showing preserved alignment across shear ranges at both time points, confirming long-term cytoskeletal adaptation. (f) eNOS expression normalized to endothelial area, demonstrating moderate or slightly increased activation under prolonged shear. Data represent mean ± SEM

Quantitative retention analysis revealed improved long-term endothelial coverage, characterized by a significant rightward shift in shear tolerance. At day 2, coverage followed a sigmoidal decline with an EC₅₀ of 33 dyn/cm² (Fig. 6d). By day 5, the EC₅₀ increased to 101 dyn/cm², indicating increased tolerance of the endothelial monolayer to applied shear. Consistent with this shift, endothelial retention at comparable shear levels was higher at day 5 than at day 2, demonstrating adaptive stabilization of the monolayer under chronic shear. This enhanced tolerance coincided with preserved junctional integrity and monolayer continuity under prolonged shear exposure. Near-complete retention was observed across forward-flow regions by day 5.

Alignment analysis further indicated persistent mechano-adaptation: orientation angles were similar between days 2 and 5 across the full shear range, with only minor deviations in transitional zones (Fig. 6e). eNOS normalized to endothelial area remained elevated at day 5 (Fig. 6f), supporting sustained nitric oxide signaling under prolonged shear.

These findings demonstrate that endothelial retention, cytoskeletal organization, and nitric oxide signaling persist over prolonged supraphysiologic shear exposure.

### Convergent shear–vorticity optima predicts endothelial retention and functional activation

To unify the structural and functional findings across geometries, we developed a multivariate regression framework relating biological responses to shear–vorticity coupling (Fig. 7a-d). Endothelial coverage was used as the reference outcome because it represents the necessary precondition for junctional integrity, cytoskeletal alignment, and downstream functional activation, including nitric oxide production. Accordingly, the shear–vorticity model relates functional readouts to the shear–vorticity range associated with preserved coverage and elevated eNOS expression and nitrite release. Coverage values were logit-transformed to enable continuous regression across the full shear spectrum.

**Fig. 7.**
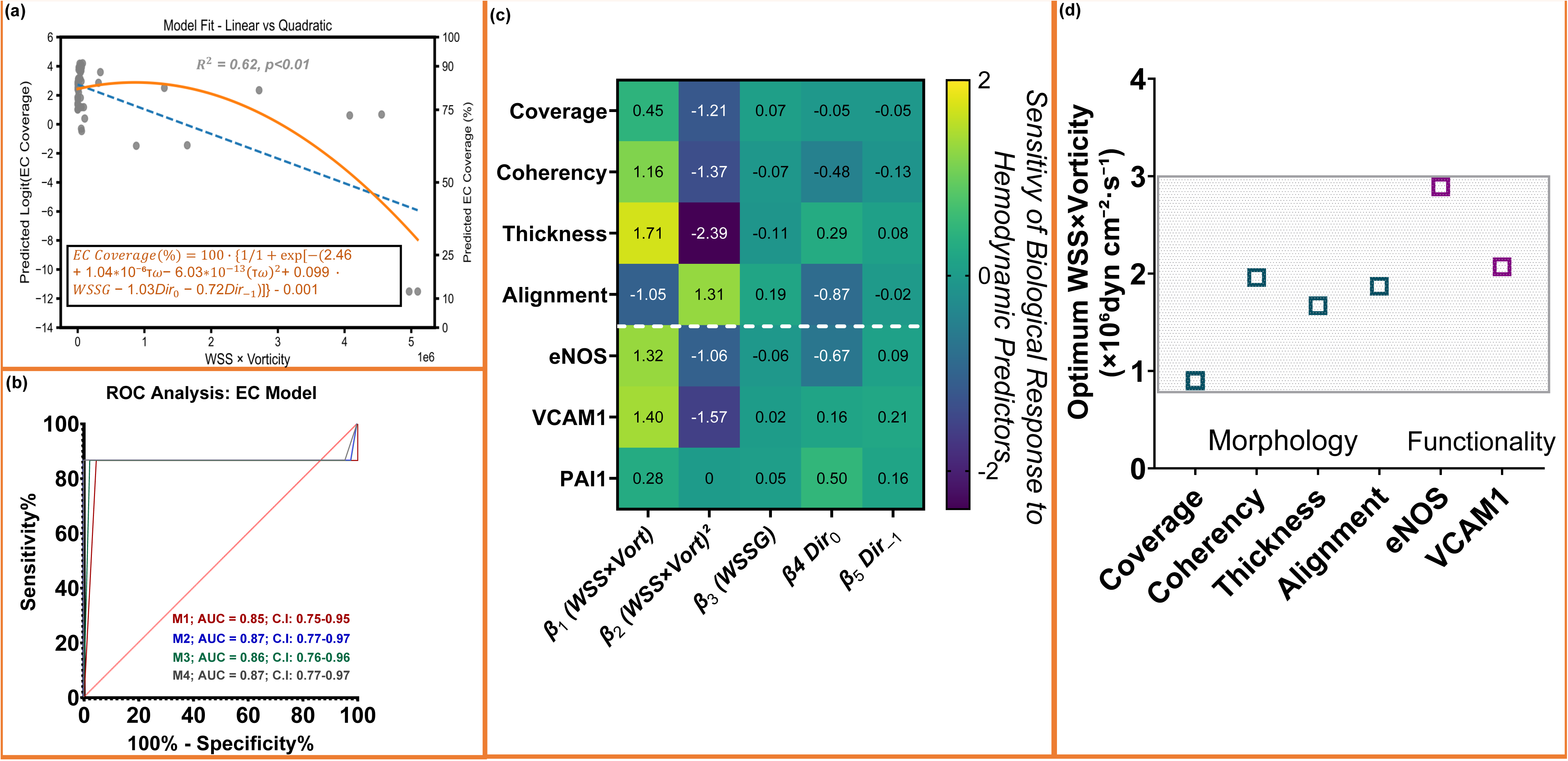
Shear-vorticity balance predicts endothelial structure, function and inflammatory tone. (a) Model comparison for endothelial coverage showing that a quadratic dependence on the combined shear–vorticity term (WSS × vorticity) provides the best fit (orange) compared with a linear model (blue). (b) ROC curves demonstrating predictive performance of shear–vorticity–based models for identifying high-retention regions (≥50% EC coverage). Adding the quadratic term yields the highest AUC (0.87), comparable to the full multivariate model incorporating WSSG and flow direction. (c) Standardized β-coefficients from multivariate regression across biological readouts. β₁ and β₂ represent the linear and quadratic components of the WSS × vorticity term; together they define the curvature and direction of each response. Positive β₁ and negative β₂ reveal a consistent parabolic dependence for coverage, coherency, junction thickness, eNOS, and VCAM-1. Alignment shows the expected sign inversion due to its directional definition, while PAI-1 is best described by a linear decline. (d) Mechanical optima (−β₁/2β₂) for each readout. Morphological features (coverage, coherency, alignment, junction thickness) converge between ≈0.9–1.9 × 10⁶ dyn·cm⁻²·s⁻¹, whereas functional responses (eNOS activation and VCAM-1 suppression) peak at ≈2.0–2.9 × 10⁶ dyn·cm⁻²·s⁻¹. The shaded region denotes the shear-vorticity regime that supports

Quadratic coupling of WSS and vorticity best predicted endothelial retention. Across four model families (linear, quadratic, exponential, logistic), the WSS × vorticity quadratic model provided the strongest performance (R² = 0.62), outperforming linear fits, while exponential/logistic models did not converge (Fig. 7a; Supplementary Table 1). The best-fit equation captured the characteristic rise-fall profile of retention with increasing shear–vorticity, with an optimum (∼0.9 × 10⁶ dyn·cm⁻²·s⁻¹) matching the experimentally observed retention peak in both 22.5° and 45° trenches. Mechanistically, the WSS × vorticity term reflects tangential shear loading superimposed with local rotational flow, capturing the interaction between shear magnitude and near-wall rotational intensity, which influences endothelial adhesion.

Because coverage transitions sharply between high-retention and low-retention regimes, we evaluated coverage using ROC analysis. A single shear–vorticity term yielded AUC = 0.85, and adding the quadratic term improved performance to AUC = 0.87, equivalent to the full multivariate model incorporating WSSG and flow direction (Fig. 7b; Supplementary Fig. 8). This confirms that the WSS × vorticity interaction is the dominant predictor of endothelial retention, with minimal additional benefit from higher-order predictors.

β-coefficient patterns further revealed a shared quadratic dependence across biological readouts. Other biological readouts were evaluated through multivariate regression (Supplementary Fig. 7a-b; Supplementary Table 2). Most outputs, including junction thickness, coherency, eNOS activation, and VCAM-1 suppression, showed the same parabolic dependence (β₁ positive, β₂ negative), indicating shared mechanical tuning. Alignment showed the expected inverse-sign pattern because angles decrease as cells rotate toward flow, while retaining a similar optimum range. PAI-1 alone was best fit by a linear model, consistent with its monotonic suppression under flow conditions associated with preserved endothelial coverage.

Optima derived from multiple biological readouts converged within a shared shear–vorticity range. Hemodynamic optima calculated from quadratic fits (–β₁/2β₂) clusteredbetween 0.8 and 2.9 × 10⁶ dyn·cm⁻²·s⁻¹ for all readouts with quadratic curvature (β₂ ≠ 0), with PAI-1 showing a linear dependence (Fig. 7d; Supplementary Table 2). Coverage showed the earliest optimum (∼0.8-0.9 × 10⁶), consistent with the threshold for detachment resistance. Cytoskeletal and junctional metrics, including coherency, alignment, and VE-cadherin junction thickness, peaked between 1.4 and 1.9 × 10⁶, consistent with increased coherency and tension-aligned actin organization. Functional outputs (eNOS activation and VCAM-1 suppression) consolidated at 2.0-2.9 × 10⁶, consistent with sustained biochemical signaling.

Together, these analyses indicate that endothelial responses across structural and functional readouts follow a shared quadratic dependence on the shear–vorticity term. Structural responses, including endothelial coverage, actin coherency, alignment, and junction thickness, whereas functional responses (eNOS activation and VCAM-1 suppression peaked toward the upper end of the same range. These findings demonstrate that coverage, cytoskeletal organization, nitric oxide production, and inflammatory marker expression are coordinated within a defined shear–vorticity, providing a predictive framework for endothelial responses under high shear.

## Discussion

Endothelial mechanobiology shapes the long-term performance of blood-contacting cardiovascular implants, yet recreating flow conditions that reinforce endothelial persistence and function remains challenging. Endothelial responses depend not only on shear magnitude but also on flow organization, integrating junctional tension, cytoskeletal alignment, nitric oxide signaling, and inflammatory signaling through coordinated VE-cadherin-actin-eNOS complexes [33,40,41], consistent with emerging evidence that maintenance of a confluent endothelial monolayer is governed by the spatial organization of near-wall flow rather than shear magnitude alone [42]. In native vessels, stable laminar shear stabilizes these structures, whereas shear below physiological thresholds or disturbed high-shear flow promotes junctional fragmentation and inflammation [43,44]. Here, we demonstrate that geometry alone can reorganize near-wall hemodynamics to approximate this balance under high shear. Mesoscale trench geometries organize near-wall flow by tuning shear and vorticity, creating spatially confined low-vorticity forward-flow niches that coordinate endothelial structure and function under supraphysiologic shear.

Across engineered geometries, endothelial cells exhibited collective behavior, with endothelial coverage extending into domains where local shear conditions alone would not be predicted to support stable adhesion. Regions characterized by elevated vorticity or abrupt shear transitions correlated with junctional thinning and reduced cytoskeletal coherency, indicating local mechanical conditions that compromise endothelial monolayer stability. In contrast, moderate shear within forward-flow domains reinforced VE-cadherin organization, increased cytoskeletal coherency, and promoted perpendicular actin alignment. These coordinated structural responses suggest that endothelial adaptation occurs collectively across the monolayer rather than at the level of isolated cells. Such behavior is consistent with collective endothelial mechano-adaptation, in which cytoskeletal forces transmitted through VE-cadherin junctions couple neighboring cells and allow local flow conditions to organize coordinated monolayer behavior [42]. Consistent with this interpretation, prior studies show that VE-cadherin junction structure reflects mechanical load distribution across endothelial monolayers and that antibody-mediated VE-cadherin blockade under flow disrupts collective alignment and structural stabilization [41,45]. Within this framework, the preservation of endothelial coverage in the 45° geometry reflects a form of collective tolerance to extreme shear, in which mechanical stresses are redistributed across the junctionally coupled monolayer rather than being concentrated at individual cell–substrate attachment sites.

Consistent with this mechanically stabilized state, the 45° geometry preserved confluent monolayers at shear stresses exceeding 200 dyn/cm² while simultaneously maintaining robust eNOS activation and nitric oxide release, consistent with the vasoprotective signaling observed across forward-flow regions. These findings indicate that endothelial tolerance to extreme shear is shaped by the combined influence of shear magnitude and rotational flow. Specifically, the preserved coverage and junctional organization observed in the 45° design suggest that geometry-optimized flow reduces denudation at the cell–material interface, allowing VE-cadherin junctions to accommodate tensile loading through the active reorganization of cellular tension forces [46], which would otherwise promote detachment on flat substrates.

Notably, endothelial coverage increased over time, indicating active shear-dependent expansion despite exposure to shear gradients beyond the predicted ideal range for endothelial function. Over five days of continuous shear exposure, the rightward shift in EC₅₀ further suggests progressive adaptation of the monolayer to sustained mechanical loading, accompanied by persistent junctional organization and nitric oxide signaling. A modest reduction in actin alignment was observed, consistent with reports that endothelial contact guidance can attenuate as monolayers mature [47]. Importantly, this change did not coincide with reductions in endothelial coverage or nitric oxide signaling.

A central advance of this work is demonstrating that structural, functional, and inflammatory endothelial responses converge within a defined shear–vorticity range. Functional activation refers specifically to eNOS expression and nitric oxide production, while inflammatory status is assessed by VCAM-1 and PAI-1 expression. Prior studies emphasized disturbed-flow metrics [48,49], but did not establish quantitative optima or demonstrate convergence across endothelial phenotypes. Here, multivariate regression revealed that retention, junction thickness, coherency, alignment, eNOS activation, and VCAM-1 responses followed a shared parabolic dependence on WSS × vorticity, while PAI-1 decreased linearly. Mechanical optima clustered between 0.8 and 2.9 × 10⁶ dyn·cm⁻²·s⁻¹ (as synthesized into a unified mechanistic schematic in Supplementary Fig. 10), spanning the transition from junctional reinforcement to maximal nitric oxide signaling and inflammatory suppression. Nitric oxide production, a functional hallmark of vasoprotective endothelium [10,50], peaked within this same regime. Beyond its role in vascular tone, NO functions as a potent anti-thrombotic mediator by suppressing platelet activation, leukocyte adhesion, and inflammatory signaling, linking the mechanically defined window to a functionally protective endothelial phenotype. This convergence establishes the shear–vorticity product as a primary design specification for blood-contacting surfaces, moving beyond descriptive metrics toward a predictive engineering framework.

Although predictive strength varied across outputs, the underlying mechanical dependence remained consistent. ROC analysis, therefore, focused on endothelial coverage, the only endpoint exhibiting a clear binary transition, enabling accurate classification of coverage monolayers based on shear–vorticity alone. Functional readouts, including eNOS expression and nitric oxide production, reflect endothelial phenotype rather than structural integrity. High endothelial coverage does not guarantee an antithrombotic state, as inflammatory or pro-coagulant phenotypes can persist within confluent monolayers and suppress NO signaling [15]. Thus, coverage serves as a classifier of structural monolayer integrity, while functional outputs provide complementary insight into endothelial phenotype within confluent monolayers.

These findings clarify longstanding inconsistencies in implant mechanobiology. Endothelial monolayers tolerate extreme shear when the flow remains steady and unidirectional [51]. Tolerance to high-magnitude shear is therefore shaped by flow organization in addition to adhesive interactions. In contrast, steep gradients or recirculation compromise monolayer integrity even at moderate shear levels. This distinction may help explain the limited durability of biochemical coatings that alter molecular interactions without addressing the local organization of near-wall flow [34]. Unlike degradable or eluting coatings, which can face challenges with late-term clinical complications in high-risk scenarios [52], geometry-encoded modulation may offer greater durability, remaining functional as long as the surface topography is preserved, and offering a potentially stable strategy for long-term mechanical circulatory support. Geometry-driven modulation of near-wall hemodynamics, therefore, represents a complementary and potentially durable strategy for hemocompatibility [32,53].

These findings have direct translational implications. Importantly, the microtrenched features were directly embossed into medical-grade UHMWPE, a clinically established implant material widely used in cardiovascular and orthopedic applications. This demonstrates that geometry-driven modulation of near-wall hemodynamics can be implemented without altering bulk material chemistry or relying on degradable or eluting coatings. Microtrenched features at the scale described here are manufacturable using established medical-grade machining, embossing, or laser texturing approaches and are compatible with materials commonly used in cardiovascular devices, including UHMWPE, PTFE, and titanium [54–57]. Because the approach relies on physical surface architecture, it integrates readily with standard hemocompatibility evaluation frameworks, including ISO 10993 and established FDA assessment pathways for blood-contacting devices [58,59]. By establishing a deterministic relationship between surface topography and endothelial response, the shear–vorticity framework provides a predictive design tool in which implant geometries can be computationally screened for residence within this regime prior to fabrication or experimental validation, enabling more efficient iterative design cycles [60].

Collectively, these findings position shear–vorticity balance as a quantifiable mechanical parameter linking surface architecture to endothelial phenotype. Importantly, the observed collective endothelial behavior extends endothelial antithrombotic phenotypes beyond the traditionally predicted “ideal” shear range, enabling endothelial persistence across both high- and low-shear domains. This expanded tolerance suggests that endothelial adaptation is governed by the spatial organization of flow rather than shear magnitude alone. By addressing the broader need to move beyond structural coverage toward functionally quiescent, antithrombotic surfaces in blood-contacting devices [61], this work provides a quantitative, predictive design rule for high-shear cardiovascular implants. The convergence across structural, functional, and inflammatory metrics further supports the robustness of this mechanobiological framework.

## Conclusion

Microtrenched UHMWPE surfaces reorganize near-wall hemodynamics into discrete shear and vorticity niches that sustain endothelial retention under supraphysiologic flow. Across trench geometries, endothelial structural integrity, nitric oxide production, and inflammatory suppression aligned within a defined WSS × vorticity window. This mechanically bounded regime provides a quantitative design rule for engineering coating-free cardiovascular surfaces that support endothelial compatibility under high-shear flow.

## Acknowledgements

This work was supported by the National Institutes of Health, National Heart, Lung, and Blood Institute (R01 HL089456 and R01 HL143247) and by the AIA Scholarship. We thank the Cornell Center for Materials Research (NSF MRSEC program) for access to profilometry instrumentation. Imaging data were acquired at the Cornell Institute of Biotechnology Imaging Facility, supported in part by NIH S10OD018516, which provides access to the shared Zeiss LSM880 confocal/multiphoton microscope. Computational resources were provided in part by the Swanson Simulation Lab. We thank Johanna De La Cruz for assistance with confocal microscopy and early imaging analysis. We also acknowledge undergraduate contributors Sarah Kazmi and Elizabeth Snavely for assistance with early image expansion and preprocessing used in the preparation of microtrench datasets.

## Author Contributions

**Aminat M Ibrahim:** Conceptualization; Methodology; Software; Investigation; Data curation; Formal analysis; Validation; Visualization; Writing – original draft. **George Zeng:** Investigation; Data curation; Writing – review & editing. **Scott J Stelick:** Resources; Methodology. **James F Antaki:** Conceptualization; Supervision; Funding acquisition; Writing – review & editing. All authors reviewed and approved the final manuscript. **Jonathan T Butcher:** Conceptualization; Supervision; Project administration; Funding acquisition; Writing – review & editing.

## Data availability

The authors declare that all data supporting the findings of this study are available within the article and its Supplementary Information. Additional source data underlying the quantitative analyses are available from the corresponding author upon reasonable request.

## References

[1] Ibrahim DM, Fomina A, Bouten CVC, Smits AIPM. Functional regeneration at the blood-biomaterial interface. Adv Drug Deliv Rev 2023;201:115085. 10.1016/J.ADDR.2023.115085.

[2] Chen MJ, Pappas GA, Massella D, Schlothauer A, Motta SE, Falk V, et al. Tailoring crystallinity for hemocompatible and durable PEEK cardiovascular implants. Biomaterials Advances 2023;146:213288. 10.1016/j.bioadv.2023.213288.

[3] Fioretta ES, Motta SE, Lintas V, Loerakker S, Parker KK, Baaijens FPT, et al. Next-generation tissue-engineered heart valves with repair, remodelling and regeneration capacity. Nature Reviews Cardiology 2020 18:2 2020;18:92–116. 10.1038/s41569-020-0422-8.

[4] Sawa S, Saito S, Morita K, Miyamoto S, Hattori M, Hino A, et al. Thirty-year outcomes of low-intensity anticoagulation for mechanical aortic valve. Heart and Vessels 2024 39:6 2024;39:549–55. 10.1007/S00380-024-02365-X.

[5] Grinstein J, Belkin MN, Kalantari S, Bourque K, Salerno C, Pinney S. Adverse Hemodynamic Consequences of Continuous Left Ventricular Mechanical Support. J Am Coll Cardiol 2023;82:70–81. 10.1016/j.jacc.2023.04.045.

[6] Wang B, Mei Z, Yang H, Gao W, Ma L, An G. Global, regional, and national burden of nonrheumatic calcific aortic valve disease based on GBD study 2021. Scientific Reports 2025 15:1 2025;15:29464-. 10.1038/s41598-025-14522-x.

[7] Smith RJ, Nasiri B, Kann J, Yergeau D, Bard JE, Swartz DD, et al. Endothelialization of arterial vascular grafts by circulating monocytes. Nature Communications 2020 11:1 2020;11:1622-. 10.1038/s41467-020-15361-2.

[8] Mirani B, Latifi N, Lecce M, Zhang X, Simmons CA. Biomaterials and biofabrication strategies for tissue-engineered heart valves. Matter 2024;7:2896–940. 10.1016/J.MATT.2024.05.036.

[9] Xiao W, Chen W, Wang Y, Zhang C, Zhang X, Zhang S, et al. Recombinant DTβ4-inspired porous 3D vascular graft enhanced antithrombogenicity and recruited circulating CD93+/CD34+ cells for endothelialization. Sci Adv 2022;8:1958. 10.1126/SCIADV.ABN1958.

[10] Cheng CK, Wang N, Wang L, Huang Y. Biophysical and Biochemical Roles of Shear Stress on Endothelium: A Revisit and New Insights. Circ Res 2025;136:752–72. 10.1161/CIRCRESAHA.124.325685.

[11] Dasi LP, Simon HA, Sucosky P, Yoganathan AP. FLUID MECHANICS OF ARTIFICIAL HEART VALVES. Clin Exp Pharmacol Physiol 2009;36:225–37. 10.1111/j.1440-1681.2008.05099.x.

[12] Gould ST, Srigunapalan S, Simmons CA, Anseth KS. Hemodynamic and cellular response feedback in calcific aortic valve disease. Circ Res 2013;113:186–97. 10.1161/CIRCRESAHA.112.300154;ISSUE:ISSUE:DOI.

[13] Gresele P, Momi S, Guglielmini G. Nitric oxide-enhancing or -releasing agents as antithrombotic drugs. Biochem Pharmacol 2019;166:300–12. 10.1016/j.bcp.2019.05.030.

[14] Peng Z, Shu B, Zhang Y, Wang M. Endothelial Response to Pathophysiological Stress. Arterioscler Thromb Vasc Biol 2019;39. 10.1161/ATVBAHA.119.312580.

[15] Chiu J-J, Chien S. Effects of Disturbed Flow on Vascular Endothelium: Pathophysiological Basis and Clinical Perspectives. Physiol Rev 2011;91:327–87. 10.1152/physrev.00047.2009.

[16] Rao J, Pan Bei H, Yang Y, Liu Y, Lin H, Zhao X. Nitric Oxide-Producing Cardiovascular Stent Coatings for Prevention of Thrombosis and Restenosis. Front Bioeng Biotechnol 2020;8:545925. 10.3389/FBIOE.2020.00578/BIBTEX.

[17] Zhu T, Gu H, Zhang H, Wang H, Xia H, Mo X, et al. Covalent grafting of PEG and heparin improves biological performance of electrospun vascular grafts for carotid artery replacement. Acta Biomater 2021;119:211–24. 10.1016/J.ACTBIO.2020.11.013.

[18] Rana D, Salave S, Benival D, Vora LK, Khunt D. Peptide-based targeting: Novel concept for thrombosis diagnosis and treatment. J Drug Deliv Sci Technol 2024;95:105612. 10.1016/j.jddst.2024.105612.

[19] Han C, Luo X, Zou D, Li J, Zhang K, Yang P, et al. Nature-inspired extracellular matrix coating produced by micro-patterned smooth muscle and endothelial cells endows cardiovascular materials with better biocompatibility. Biomater Sci 2019;7:2686–701. 10.1039/C9BM00128J.

[20] Haritz JL, Pflaum M, Güntner HJ, Katsirntaki K, Hegermann J, Hehnen F, et al. Citrate-Coated Iron Oxide Nanoparticles Facilitate Endothelialization of Left Ventricular Assist Device Impeller for Improved Antithrombogenicity. Advanced Science 2025;12. 10.1002/advs.202408976.

[21] Zhou K, Li Y, Zhang L, Jin L, Yuan F, Tan J, et al. Nano-micrometer surface roughness gradients reveal topographical influences on differentiating responses of vascular cells on biodegradable magnesium. Bioact Mater 2021;6:262–72. 10.1016/j.bioactmat.2020.08.004.

[22] Liu S, Yan J, Gao M, Yang H. Research progress in the regulation of endothelial cells and smooth muscle cells using a micro–nanostructure. Biomed Eng Online 2025;24:6. 10.1186/s12938-025-01337-0.

[23] Robotti F, Franco D, Bänninger L, Wyler J, Starck CT, Falk V, et al. The influence of surface micro-structure on endothelialization under supraphysiological wall shear stress. Biomaterials 2014;35:8479–86. 10.1016/j.biomaterials.2014.06.046.

[24] Wilson AC, Neuenschwander PF, Chou S-F. Engineering Approaches to Prevent Blood Clotting from Medical Implants. Arch Biomed Eng Biotechnol 2019;1:.000510. 10.33552/ABEB.2018.01.000510.

[25] Lancellotti P, Aqil A, Musumeci L, Jacques N, Ditkowski B, Debuisson M, et al. Bioactive surface coating for preventing mechanical heart valve thrombosis. Journal of Thrombosis and Haemostasis 2023;21:2485–98. 10.1016/j.jtha.2023.05.004.

[26] Zhao J, Feng Y. Surface Engineering of Cardiovascular Devices for Improved Hemocompatibility and Rapid Endothelialization. Adv Healthc Mater 2020;9:2000920. 10.1002/ADHM.202000920;PAGEGROUP:STRING:PUBLICATION.

[27] Haritz JL, Pflaum M, Güntner HJ, Katsirntaki K, Hegermann J, Hehnen F, et al. Citrate-Coated Iron Oxide Nanoparticles Facilitate Endothelialization of Left Ventricular Assist Device Impeller for Improved Antithrombogenicity. Advanced Science 2025;12. 10.1002/advs.202408976.

[28] Clark AG, Maitra A, Jacques C, Bergert M, Pérez-González C, Simon A, et al. Self-generated gradients steer collective migration on viscoelastic collagen networks. Nature Materials 2022 21:10 2022;21:1200–10. 10.1038/s41563-022-01259-5.

[29] Dessalles CA, Leclech C, Castagnino A, Barakat AI. Integration of substrate- and flow-derived stresses in endothelial cell mechanobiology. Commun Biol 2021;4:764. 10.1038/s42003-021-02285-w.

[30] Mannino RG, Myers DR, Ahn B, Wang Y, Margo Rollins, Gole H, et al. “Do-it-yourself in vitro vasculature that recapitulates in vivo geometries for investigating endothelial-blood cell interactions.” Scientific Reports 2015 5:1 2015;5:12401-. 10.1038/srep12401.

[31] Wei B, Chen L, Huang X, Chi F, Li G, Yang L, et al. Microenvironment-Responsive Biomimetic Bioprosthetic Valve with Antithrombosis and Immunoregulation Performance. ACS Appl Mater Interfaces 2025;17:18160–78. 10.1021/acsami.5c01314.

[32] Conner AA, David D, Yim EKF. The Effects of Biomimetic Surface Topography on Vascular Cells: Implications for Vascular Conduits. Adv Healthc Mater 2024;13:2400335. 10.1002/ADHM.202400335.

[33] Dolan JM, Meng H, Singh S, Paluch R, Kolega J. High Fluid Shear Stress and Spatial Shear Stress Gradients Affect Endothelial Proliferation, Survival, and Alignment. Ann Biomed Eng 2011;39:1620. 10.1007/S10439-011-0267-8.

[34] Frendl CM, Tucker SM, Khan NA, Esch MB, Kanduru S, Cao TM, et al. Endothelial retention and phenotype on carbonized cardiovascular implant surfaces. Biomaterials 2014;35:7714–23. 10.1016/J.BIOMATERIALS.2014.05.075.

[35] Zhong T, Li Y, He X, Liu Y, Dong Y, Ma H, et al. Adaptation of endothelial cells to shear stress under atheroprone conditions by modulating internalization of vascular endothelial cadherin and vinculin. Ann Transl Med 2020;8:1423–1423. 10.21037/atm-20-3426.

[36] He W, Ibrahim AM, Karmakar A, Tuli S, Butcher JT, Antaki JF. Computational Fluid Dynamic Optimization of Micropatterned Surfaces: Towards Biofunctionalization of Artificial Organs. Bioengineering 2024;11:1092. 10.3390/BIOENGINEERING11111092/S1.

[37] Gould RA, Butcher JT. Isolation of valvular endothelial cells. J Vis Exp 2010. 10.3791/2158.

[38] Brezovjakova H, Tomlinson C, Mohd-Naim N, Swiatlowska P, Erasmus JE, Huveneers S, et al. Junction Mapper is a novel computer vision tool to decipher cell–cell contact phenotypes. Elife 2019;8:e45413. 10.7554/ELIFE.45413.

[39] Giese W, Albrecht JP, Oppenheim O, Akmeriç EB, Kraxner J, Schmidt D, et al. Polarity-JaM: an image analysis toolbox for cell polarity, junction and morphology quantification. Nature Communications 2025 16:1 2025;16:1474-. 10.1038/s41467-025-56643-x.

[40] Roux E, Bougaran P, Dufourcq P, Couffinhal T. Fluid Shear Stress Sensing by the Endothelial Layer. Front Physiol 2020;11:533349. 10.3389/FPHYS.2020.00861/BIBTEX.

[41] Cronin NM, Dawson LW, DeMali KA. Shear stress-stimulated AMPK couples endothelial cell mechanics, metabolism and vasodilation. J Cell Sci 2024;137. 10.1242/jcs.262232.

[42] Leclech C, Gonzalez-Rodriguez D, Villedieu A, Lok T, Déplanche AM, Barakat AI. Topography-induced large-scale antiparallel collective migration in vascular endothelium. Nat Commun 2022;13. 10.1038/s41467-022-30488-0.

[43] Suriyagandhi V, Ma Y, Paparozzi V, Guarnieri T, Di Pietro B, Dimitri GM, et al. Mechanotransduction and inflammation: An updated comprehensive representation. Mechanobiology in Medicine 2024;3:100112. 10.1016/J.MBM.2024.100112.

[44] Harrison DG, Widder J, Grumbach I, Chen W, Weber M, Searles C. Endothelial mechanotransduction, nitric oxide and vascular inflammation. J Intern Med 2006;259:351–63. 10.1111/J.1365-2796.2006.01621.X.

[45] Wang W, Lollis EM, Bordeleau F, Reinhart-King CA. Matrix stiffness regulates vascular integrity through focal adhesion kinase activity. The FASEB Journal 2019;33:1199–208. 10.1096/fj.201800841R.

[46] Chandurkar MK, Mittal N, Royer-Weeden SP, Lehmann SD, Michels EB, Haarman SE, et al. Transient low shear-stress preconditioning influences long-term endothelial traction and alignment under high shear flow. American Journal of Physiology-Heart and Circulatory Physiology 2024;326:H1180–92. 10.1152/ajpheart.00067.2024.

[47] Leclech C, Krishnamurthy A, Muller L, Barakat AI. Distinct Contact Guidance Mechanisms in Single Endothelial Cells and in Monolayers. Adv Mater Interfaces 2023;10. 10.1002/admi.202202421.

[48] Driscoll K, Cruz AD, Butcher JT. Inflammatory and Biomechanical Drivers of Endothelial-Interstitial Interactions in Calcific Aortic Valve Disease. Circ Res 2021;128:1344–70. 10.1161/CIRCRESAHA.121.318011;PAGE:STRING:ARTICLE/CHAPTER.

[49] Baeyens N, Bandyopadhyay C, Coon BG, Yun S, Schwartz MA. Endothelial fluid shear stress sensing in vascular health and disease. J Clin Invest 2016;126:821–8. 10.1172/JCI83083.

[50] Loscalzo J. Nitric oxide in vascular biology: elegance in complexity. Journal of Clinical Investigation 2024;134. 10.1172/JCI176747.

[51] Lim XR, Harraz OF. Mechanosensing by Vascular Endothelium. Annu Rev Physiol 2024;86:71–97. 10.1146/annurev-physiol-042022-030946.

[52] Jang WJ, Park IH, Oh JH, Choi KH, Song Y Bin, Hahn J-Y, et al. Efficacy and safety of durable versus biodegradable polymer drug-eluting stents in patients with acute myocardial infarction complicated by cardiogenic shock. Sci Rep 2024;14:6301. 10.1038/s41598-024-56925-2.

[53] Łopianiak I, Wojasiński M, Kuźmińska A, Trzaskowska P, Butruk-Raszeja BA. The effect of surface morphology on endothelial and smooth muscle cells growth on blow-spun fibrous scaffolds. Journal of Biological Engineering 2021 15:1 2021;15:27-. 10.1186/S13036-021-00278-1.

[54] Hussain M, Naqvi RA, Abbas N, Khan SM, Nawaz S, Hussain A, et al. Ultra-High-Molecular-Weight-Polyethylene (UHMWPE) as a Promising Polymer Material for Biomedical Applications: A Concise Review. Polymers (Basel) 2020;12:323. 10.3390/polym12020323.

[55] Kumar R, Rezapourian M, Rahmani R, Maurya HS, Kamboj N, Hussainova I. Bioinspired and Multifunctional Tribological Materials for Sliding, Erosive, Machining, and Energy-Absorbing Conditions: A Review. Biomimetics 2024;9:209. 10.3390/biomimetics9040209.

[56] Schnell G, Staehlke S, Duenow U, Nebe JB, Seitz H. Femtosecond Laser Nano/Micro Textured Ti6Al4V Surfaces—Effect on Wetting and MG-63 Cell Adhesion. Materials 2019;12:2210. 10.3390/ma12132210.

[57] Ionescu AC, Brambilla E, Azzola F, Ottobelli M, Pellegrini G, Francetti LA. Laser microtextured titanium implant surfaces reduce in vitro and in situ oral biofilm formation. PLoS One 2018;13:e0202262. 10.1371/journal.pone.0202262.

[58] Weber M, Steinle H, Golombek S, Hann L, Schlensak C, Wendel HP, et al. Blood-Contacting Biomaterials: In Vitro Evaluation of the Hemocompatibility. Front Bioeng Biotechnol 2018;6:395774. 10.3389/FBIOE.2018.00099/EPUB.

[59] Use of International Standard ISO 10993-1, “Biological evaluation of medical devices-Part 1: Evaluation and testing within a risk management process” Guidance for Industry and Food and Drug Administration Staff Preface Public Comment 2023.

[60] Jneid H, Chikwe J, Arnold S V., Bonow RO, Bradley SM, Chen EP, et al. 2024 ACC/AHA Clinical Performance and Quality Measures for Adults With Valvular and Structural Heart Disease. J Am Coll Cardiol 2024;83:1579–613. 10.1016/j.jacc.2023.12.006.

[61] Oubari H, Berkane Y, Jeljeli M, Lellouch AG, Smadja DM. Engineering the Future of Stem Cells in Vascular Reconstruction: A Leap Towards Functional Endothelialized Tissue-Engineered Vascular Conduits. Stem Cell Rev Rep 2025;21:2796–806. 10.1007/s12015-025-10968-8.

