## Supplemental Figure 1-10, Supplemental Table 1-4 for "Geometry–Encoded Microtrenches Stabilize Endothelium on High Shear Biomaterial Surfaces"

### Supplementary information

#### Supplemental Methods

The procedures for generating endothelial coverage, VE-cadherin thickness, actin orientation, and marker expression masks are described in the Main Methods. This section provides additional validation steps and quality control analyses that ensure the accuracy and reproducibility of these measurements.

##### Endothelial Coverage and Mask Validation

To confirm that thresholded masks reflected true endothelial boundaries, each coverage mask was compared directly to the underlying VE-cadherin and phalloidin channels. Spatial colocalization was assessed using the JACoP plugin in Fiji for pixel overlap coefficient and Manders' M2 coefficient, which quantify the fraction of biological signal captured by the mask. Across all images, the overlap coefficient exceeded 0.90, and Manders M2 averaged 0.98, indicating excellent agreement between masks and raw signal. Representative overlays and the colocalization values are shown in Supplementary Fig. 6.

##### VE-Cadherin Junction Thickness Quality Control

For junctional thickness measurements, further precision was achieved by enabling sub-pixel sampling in the ImageJ profile plotting tool. This allowed thickness values to be extracted with sub-micron accuracy. Junctions were also examined in upstream and downstream orientations within the same cell, enabling paired comparisons across shear domains. Regions containing disrupted or discontinuous junctions were excluded from analysis. Supplementary Fig. 9 provides representative junction profiles and example ROIs.

##### Actin Orientation and Coherency Outputs

OrientationJ generates both mean orientation values and full angular distributions. While the Main Methods present mean values and coherency indices, complete distributions for each region, detailing the range and frequency of actin fiber angles, are provided in Extended Data Figs. 3 and 4. These distributions confirm that regions classified as highly coherent indeed exhibit narrow angular spreads, whereas disturbed-flow regions show broader distributions and reduced order.

##### Nuclear Polarity ROI Annotation

Extended examples of nuclear polarity measurements, including annotated cell and nuclear centroids within upstream, base, and downstream regions, are shown in Supplementary Fig. 5. These annotations illustrate how the coordinate axes were assigned relative to the flow direction and verify consistent application of ROI boundaries across all images. Nuclear polarity was quantified by measuring the displacement of the nuclear centroid relative to the cell centroid along the flow axis. For all datasets, the coordinate origin (y = 0) was defined at the channel inlet, with y increasing in the downstream direction. The polarity index was computed as:

$$Polarity\ Index = \frac{y_{nuc} - y_{cell}}{Major\ axis}$$

which normalizes centroid displacement to cell size. Negative values indicate upstream nuclear positioning (against the direction of flow), the canonical mechanosensory response of endothelial cells under laminar shear, whereas positive values represent downstream displacement.

##### Computational Region Mapping

CFD-derived mechanical metrics were mapped directly onto the biological images to ensure regional correspondence between flow features and endothelial responses. For each trench geometry, Fluent-exported fields for wall shear stress (WSS), vorticity, wall-shear-stress gradient (WSSG), and flow direction were saved as spatial coordinate masks defining the upstream, base, and downstream regions. These masks were registered to confocal images using the trench contours as fiducials, enabling each microscopic field of view to be assigned the exact local mechanical values predicted by the CFD model.

Mesh characteristics for all simulations are reported in Supplementary Table 4. Each geometry was meshed with fine near-wall refinement to resolve steep gradients at ridge peaks and within recirculation pockets. Total element counts of ~$1.6\times10^5$ to $3.0\times10^5$, depending on geometry, with controlled aspect ratios (< 16). The resulting solutions produced smooth, numerically stable WSS and vorticity fields suitable for downstream biological mapping.

##### Modeling Frameworks and Mathematical Derivations

Biological outputs were modeled as functions of shear and vorticity. Because coverage endpoints approached bounds at 0% or 100%, the values were transformed prior to regression using a standard logit function with a small offset to avoid singularities. The transformation is expressed as:

$$Logit\ (y) = \ ln\ (\frac{y + \epsilon}{1 - (y + \epsilon)})$$

Where $\epsilon = 0.001$.

Multivariate models were fit as linear or quadratic functions of WSS, vorticity, and their interaction. For quadratic models of the form:

$$y = \ \beta_0 + \ \beta_1 x_1 + \beta_2 x_2 + \cdots$$

the mechanical optimum was determined by differentiating with respect to X and solving the turning point:

$$X_{opt} = -\frac{\beta_1}{2\beta_2}$$

This calculation yielded the shear–vorticity values maximizing each biological output. All regression coefficients, model fits, and derived optima are reported in Supplementary Table 2, with consolidated results presented in Fig. 7d.

Binary endothelial retention (≥50%) was evaluated using ROC analysis. ROC curves and AUC values for individual predictors (WSS, vorticity, WSSG, flow direction) and combined models appear in Supplementary Fig. 8.

For computational use, the regression coefficients ($\beta_1$ and $\beta_2$) provide a compact rule set. $\beta_1$ reflects initial sensitivity to shear–vorticity loading, $\beta_2$ defines response curvature, and $-\beta_1/2\beta_2$ yields the mechanical optimum. These coefficients allow CFD modelers to map WSS and vorticity fields directly onto biological outcomes, bridging flow simulation and predicted endothelial behavior. All regression coefficients, model fits, and derived optima are reported in Supplementary Table 2, with consolidated results presented in Fig. 7d.

##### Data Handling and Replication

Experiments were performed across independent biological replicates, where each replicate corresponds to a separately fabricated and endothelialized microtrench sample. Biological replicate numbers ranged from 5 to 8 across assays and geometries, with exact n reported in the corresponding figure legends.
Each biological replicate was imaged in two to three technical replicates, yielding multiple fields of view per region and ensuring adequate sampling depth for all quantitative analyses. OrientationJ results were cross-validated using whole-cell alignment derived from VE-cadherin masks. Outlier exclusion was limited to non-biological imaging artifacts, such as edge-of-field dropout or incomplete staining.

#### Supplementary Information Figures

Supplementary Fig 1: Pressure and flow classification within microtrenches.
Supplementary Fig 2: Geometry-based segmentation of microtrenches for local shear mapping.
Supplementary Fig 3: Cytoskeletal coherency and intra-region junctional behavior across shear niches.
Supplementary Fig 4: Polarity and intra-region endothelial responses across shear domains.
Supplementary Fig. 5: Polarity index definition.
Supplementary Fig. 6: Validation of endothelial coverage masks using colocalization metrics.

Supplementary Fig. 7: Quadratic vs. linear modeling of endothelial responses to shear–vorticity balance.
Supplementary Fig. 8: ROC-based comparison of shear–vorticity models for EC coverage.
Supplementary Fig. 9: Representative VE-Cadherin Junction Thickness Measurement.
Supplementary Fig. 10: Unified mechanistic framework describing endothelial adaptation across shear–vorticity regimes

#### Supplemental Tables

Supplementary Table 1: Summary of model performance across biological outputs
Supplementary Table 2: Regression coefficients and mechanical optima derived from quadratic fits
Supplementary Table 3: Antibody List.
Supplementary Table 4: CFD Mesh Resolution Parameters

#### Supplemental Movies

Supplementary Movie a: 0° trench showing endothelial retention after 48h of shear
Supplementary Movie b: 22.5° trench showing endothelial retention after 48h of shear
Supplementary Movie c: 45° trench showing endothelial retention after 48h of shear

### Supplementary Figures

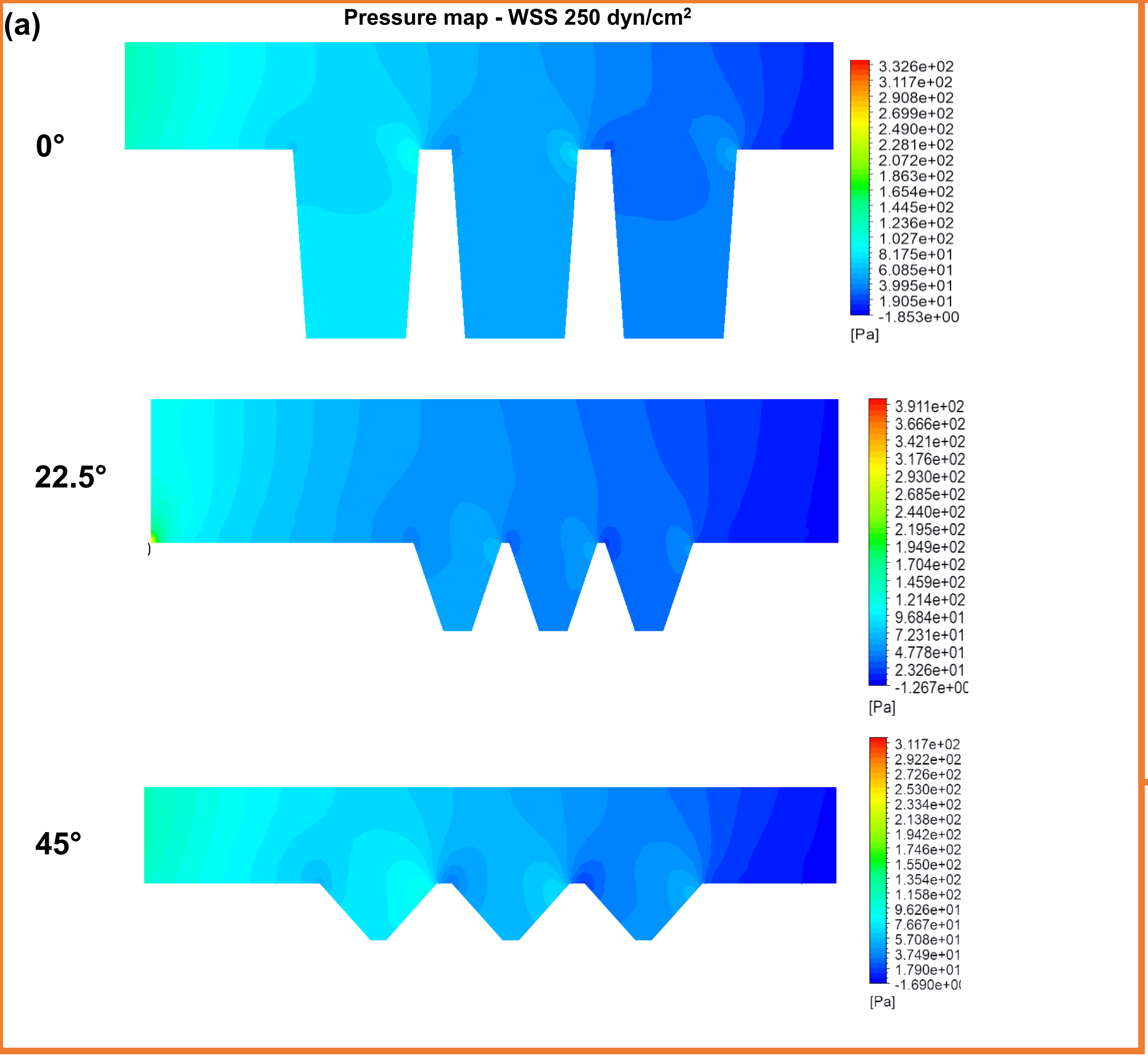


**(b)** **Flow classification parameters**

| **Parameters** | **Classification Strategy** | | |
|---|---|---|---|
| **Wall Shear Stress (WSS)** | ↓ Thromboprone (<10 dyn/cm2) | = EC Ideal (10-43 dyn/cm2) | ↑ High shear (>43 dyn/cm2) |
| **Vorticity** | ↓ Low | = Intermediate | ↑ High |
| **Flow direction** | = Neutral | → Positive = Forward | ← Negative = Recirculating |
| **Wall shear stress gradient (WSSG)** | = Stagnant | → Positive = Forward | ← Negative |

**Supplementary Fig. 1 | Pressure and flow classification within microtrenches.** (a) Pressure distribution maps reveal geometry-dependent gradients, with elevated pressure near recirculating zones and lower pressure within forward-flow regions.(b) Summary matrix classifying flow domains as thrombo-prone (<10 dyn cm$^{-2}$), endothelial-ideal (10–43 dyn cm$^{-2}$), or high-shear (>43 dyn cm$^{-2}$), based on corresponding vorticity, flow direction, and wall shear stress gradients. These parameters collectively define the engineered microenvironments used to modulate endothelial collective behavior in subsequent assays.

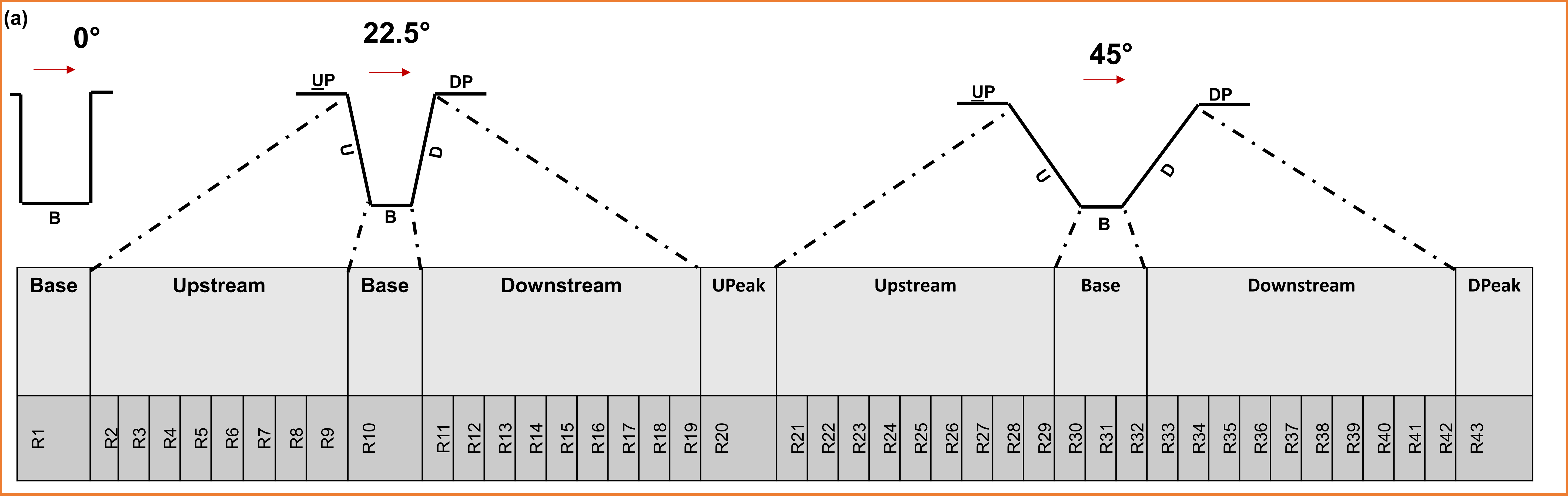


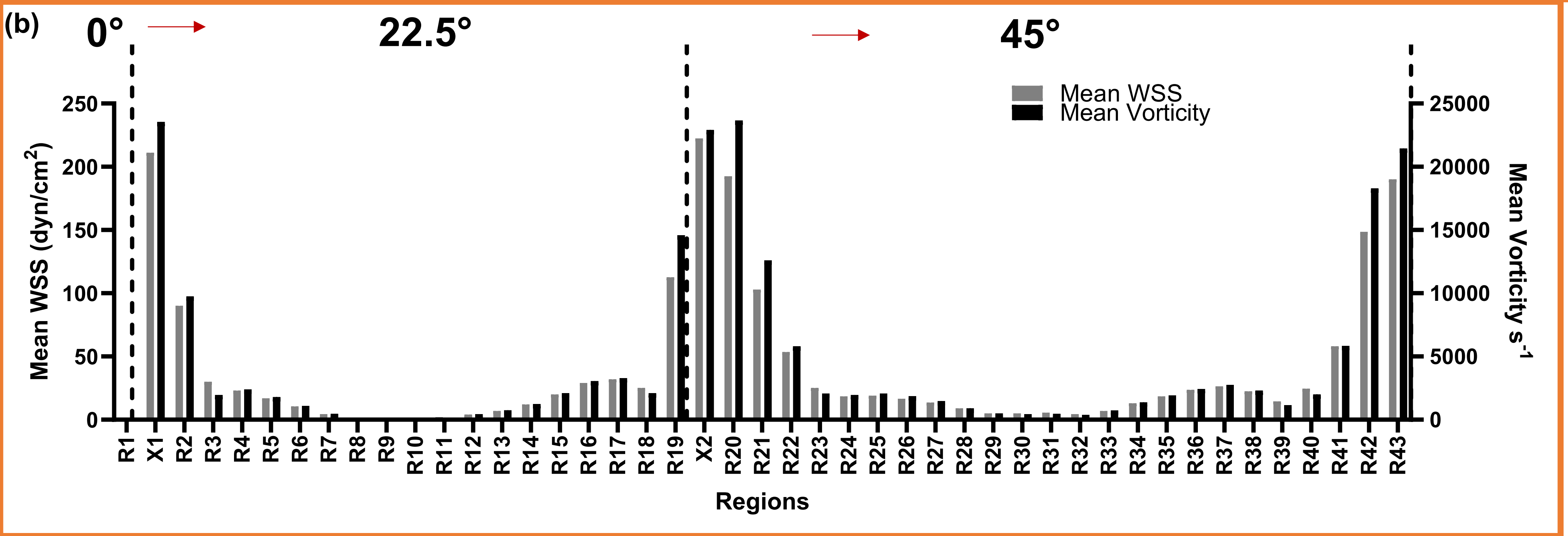


**Supplementary Fig. 2 | Geometry-based segmentation of microtrenches for local shear mapping**. (a). Schematic of geometry-based regional classification 0°, 22.5°, 45° showing upstream, base, and downstream sub-regions (R1–R43). (b) Spatial distribution of mean wall shear stress (WSS) and mean vorticity across all regions. Peaks correspond to ridge acceleration zones, while valleys denote basal recirculation pockets. These quantitative profiles illustrate how geometry defines the shear landscape used for biological analysis.

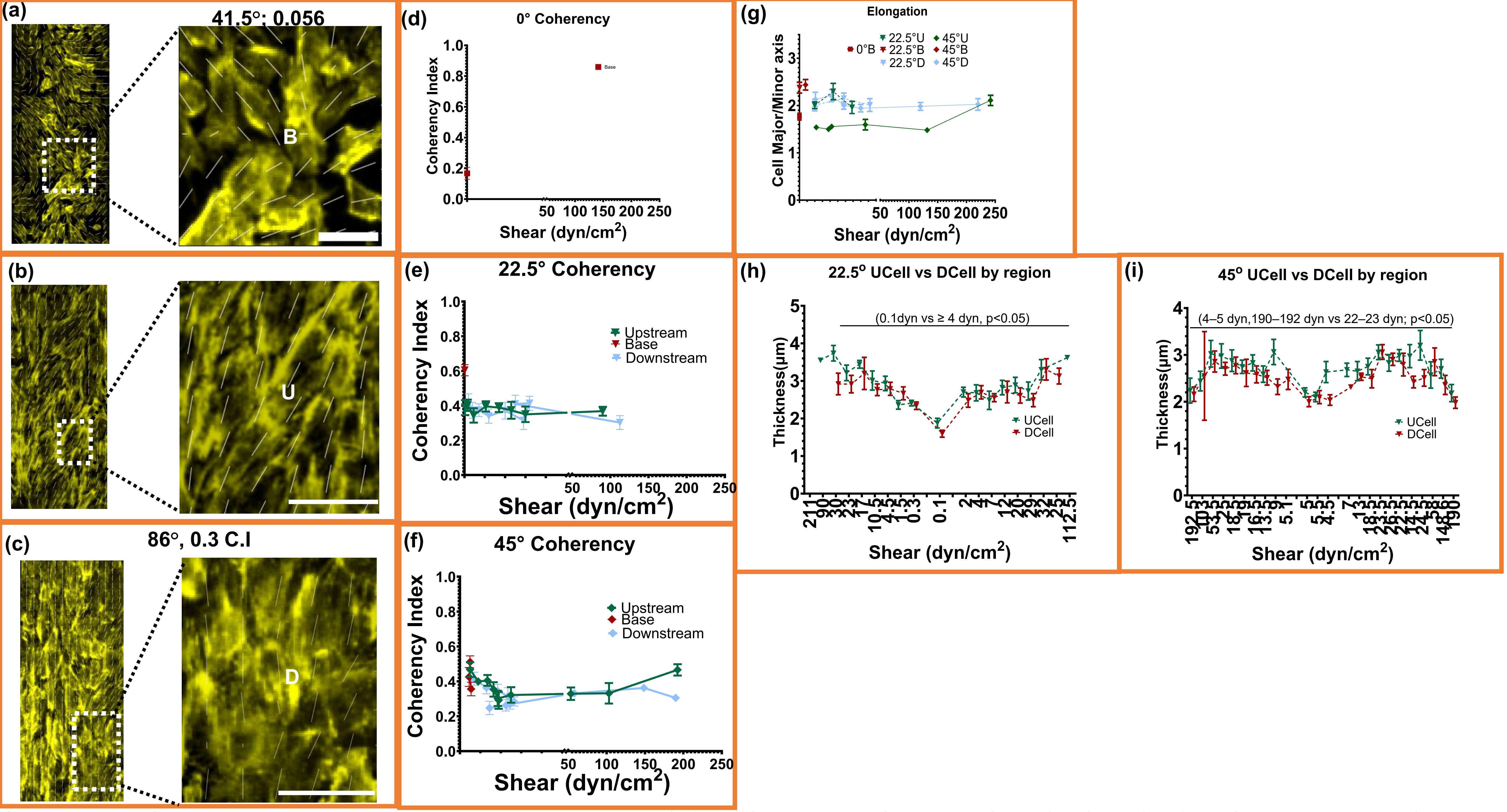


**Supplementary Fig. 3 | Cytoskeletal coherency and intra-region junctional behavior across shear niches.** (a–c) Immunofluorescence images of endothelial actin filaments (yellow) within 0°, 22.5°, and 45° microtrenches showing local fiber orientation (white lines), coherency index (C.I.), and mean orientation angle. Insets highlight representative Upstream (U), Base (B), and Downstream (D) regions. (d–f) Quantification of actin coherency across shear ranges. Coherency was lowest in the 0° base (C.I. < 0.2) under near-stagnant flow. In 22.5° and 45° geometries, coherency increased to ≈0.25–0.4 under forward shear, indicating partial but reproducible alignment of stress fibers. Local reductions within high-vorticity or recirculating zones reflect disrupted fiber continuity. (g) Cell elongation (aspect ratio) across shear levels; no geometry-dependent differences detected. (h–i) Comparison of upstream-edge (UCell) and downstream-edge (DCell) VE-cadherin junction thickness within shear-matched subregions in 22.5° and 45° geometries, showing that variation arises from geometry-driven shear rather than positional polarity. Data represent mean ± SEM; n = 8 biological replicates with triplicate measurements per sample. One-way ANOVA with Tukey's HSD; Pearson correlation. **** $p < 0.0001$, *** $p < 0.0005$, ** $p < 0.005$, * $p < 0.05$, ns = not significant.

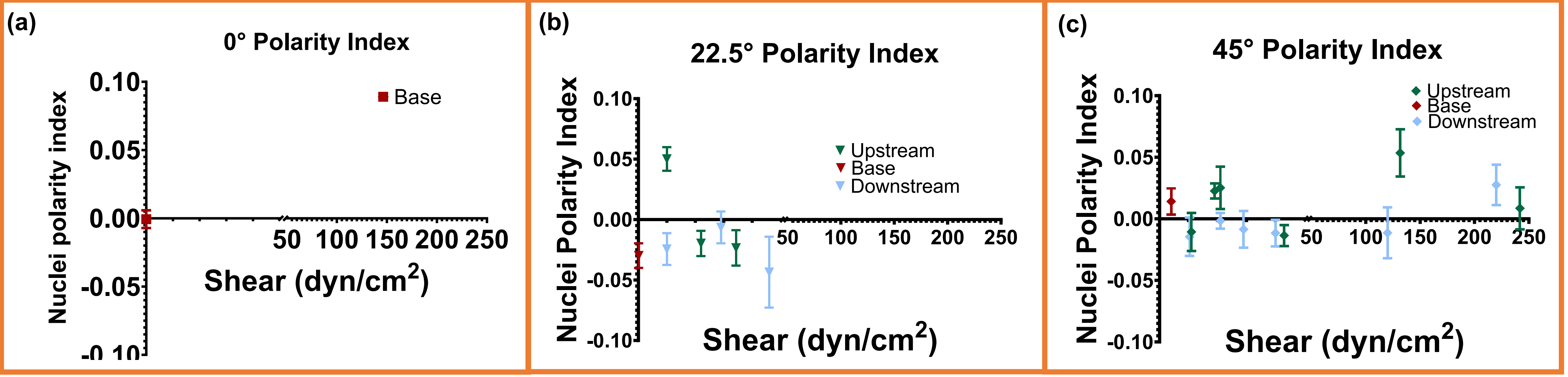


**Supplementary Fig. 4 | Polarity and intra-region endothelial responses across shear domains.** (a-c)Nuclear polarity as a function of local shear stress for 0°, 22.5°, and 45° geometries. Upstream nuclear bias emerged consistently within stable forward-flow zones, while near-stagnant and reversal regions showed variable orientation. These polarity trends parallel the shear-dependent junctional and cytoskeletal remodeling reported in Fig. 3. Data represent mean ± SEM; n = 8 biological samples.

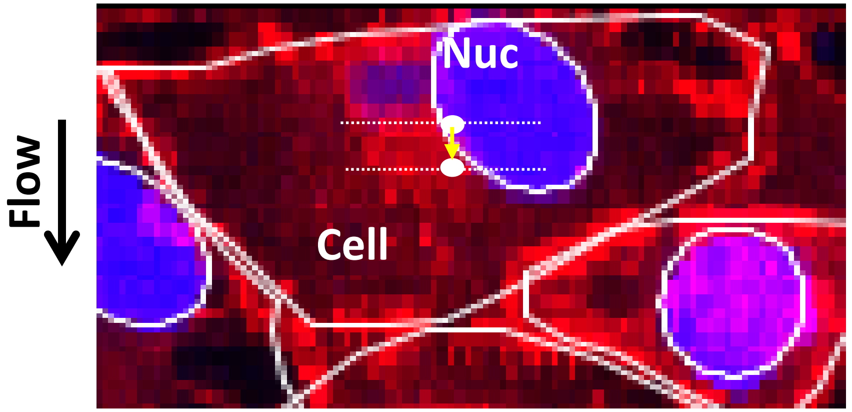


$$Polarity\ Index = \frac{pflow}{Cell\ Major\ Axis};\ \text{where}$$

$$p_{flow}(dX) = X_{nucleus\ centroid} - X_{cell\ centroid}$$

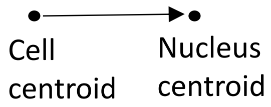


**Supplementary Fig. 5 | Polarity index definition.** Schematic illustrating computation of the nuclear polarity index based on displacement of the nuclear centroid relative to the cell centroid along the flow axis. Negative values indicate upstream polarization.

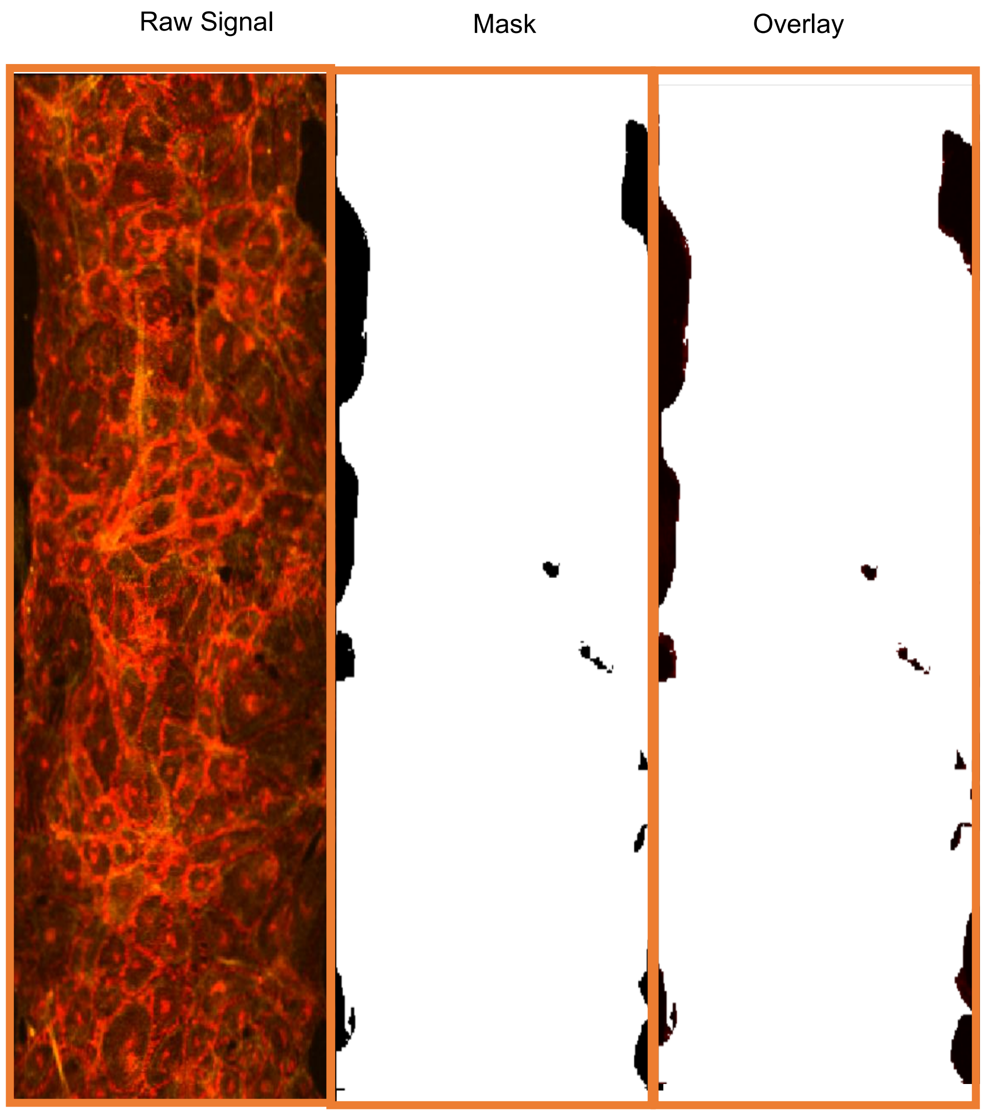


| Manders M1 | Manders M2 | Overlap Coefficient |
|---|---|---|
| 1 | 0.98 | 0.9 |

**Supplementary Fig. 6 | Validation of endothelial coverage masks using colocalization metrics.** Representative example of raw ACTIN/VE-cadherin signal, corresponding binary mask, and signal–mask overlay. Mask fidelity was evaluated using JACoP-derived metrics, including Manders' M1 and M2 coefficients and pixel overlap coefficient. Across all images, Manders' M2 exceeded 0.98 and overlap coefficients exceeded 0.90, confirming accurate inclusion of endothelial signal within the mask.

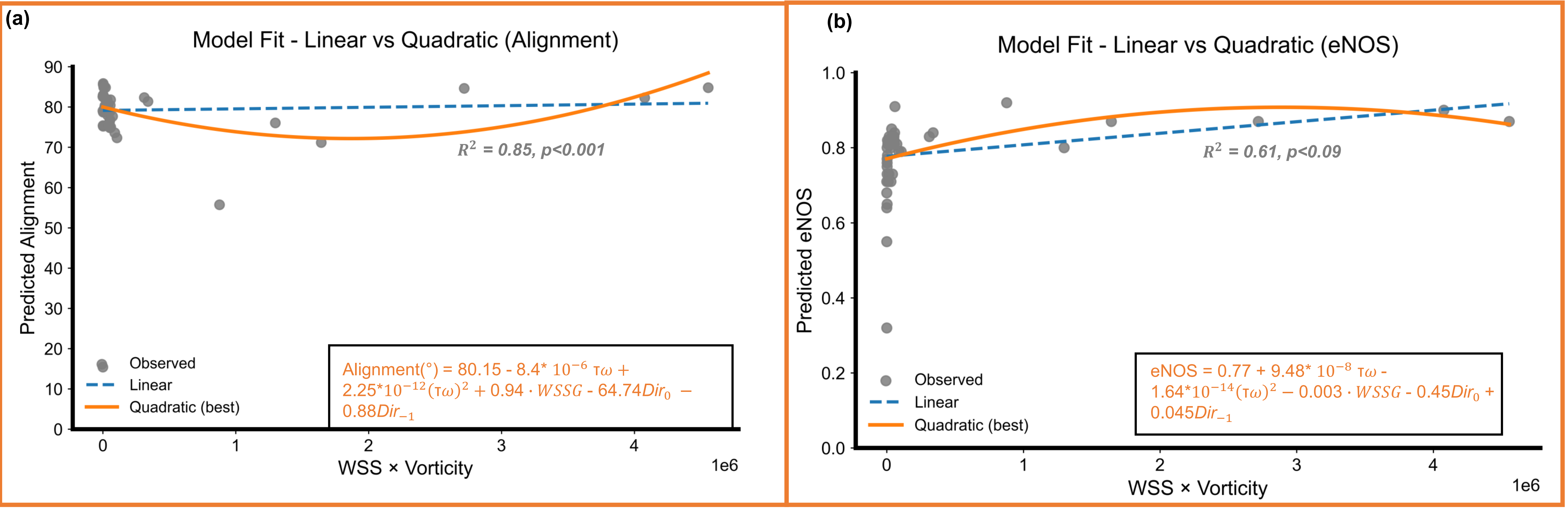


**Supplementary Fig. 7 | Quadratic vs. linear modeling of endothelial responses to shear–vorticity balance.** Representative fits for (a) alignment and (b) eNOS, demonstrating superior performance of quadratic models relative to linear models.

(a)

| Model | Predictors | AUC | Best cutoff | Sensitivity | Specificity | LR | Youden J | Note |
|---|---|---|---|---|---|---|---|---|
| | EC Coverage | | | | | | | |
| 1 | β0 + β1(WSS×Vort) | 0.8474 | > 0.9995 | 86.67 | 95.56 | 19.5 | 181.23 | Strong basic predictor |
| 2 | β0 + β1$(WSS \times Vort)^2$ | 0.8681 | > 0.9740 | 86.67 | 100 | | 185.67 | Improves prediction slightly |
| 3 | β0 + β1$(WSS \times Vort)^2$ +WSSG | 0.8585 | > 0.9710 | 86.67 | 97.78 | 39 | 183.45 | No improvement |
| 4 | β0 + β1$(WSS \times Vort)^2$ +WSSG+Dir0+Dir-1 | 0.8696 | > 0.9800 | 86.67 | 100 | | 185.67 | BEST |

**Supplementary Fig. 8 | ROC-based comparison of shear–vorticity models for EC coverage.** ROC curves comparing model performance with WSS × vorticity alone, with quadratic term, with WSSG added, and using the full multivariate model. Quadratic shear–vorticity achieved optimal sensitivity (86.7%) and specificity (95.6–100%), confirming dominance of the nonlinear mechanical term.

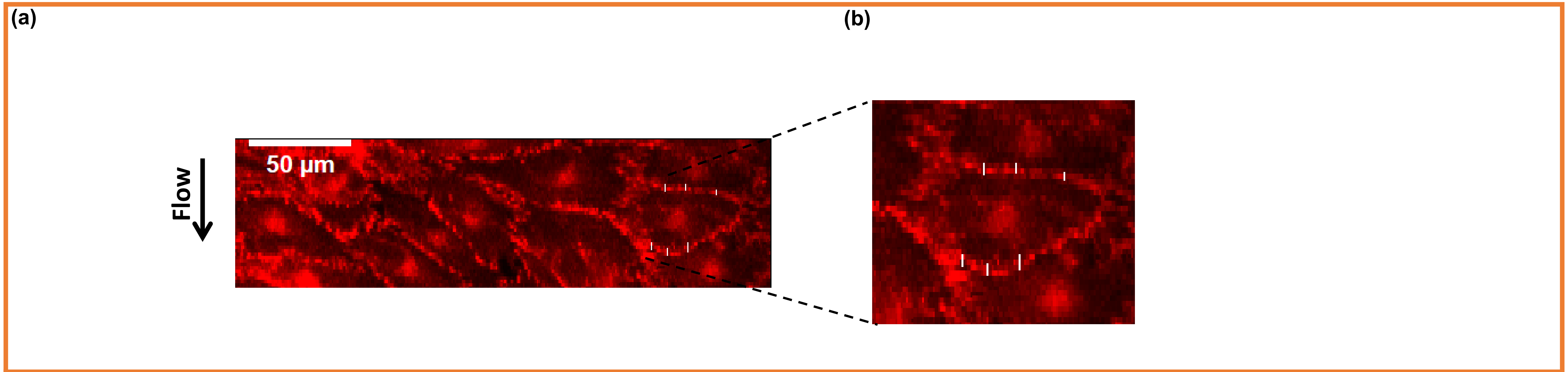


(c)

| | Cells: | |
|---|---|---|
| | U1 | D1 |
| | 4.982 | 4.53 |
| | 3.963 | 1.415 |
| | 5.096 | 2.548 |
| Mean | **4.68** | **2.831** |

**Supplementary Fig. 9 | Representative VE-Cadherin Junction Thickness Measurement.** (a) Raw VE-cadherin immunofluorescence from microtrenched monolayers. (b) Enlarged view showing perpendicular line ROIs used to measure junction thickness across continuous junction segments. (c) Example of quantified junction width values for upstream (U1) and downstream (D1) cells. These images illustrate the morphology and measurement approach used for quantitative analysis across shear niches.

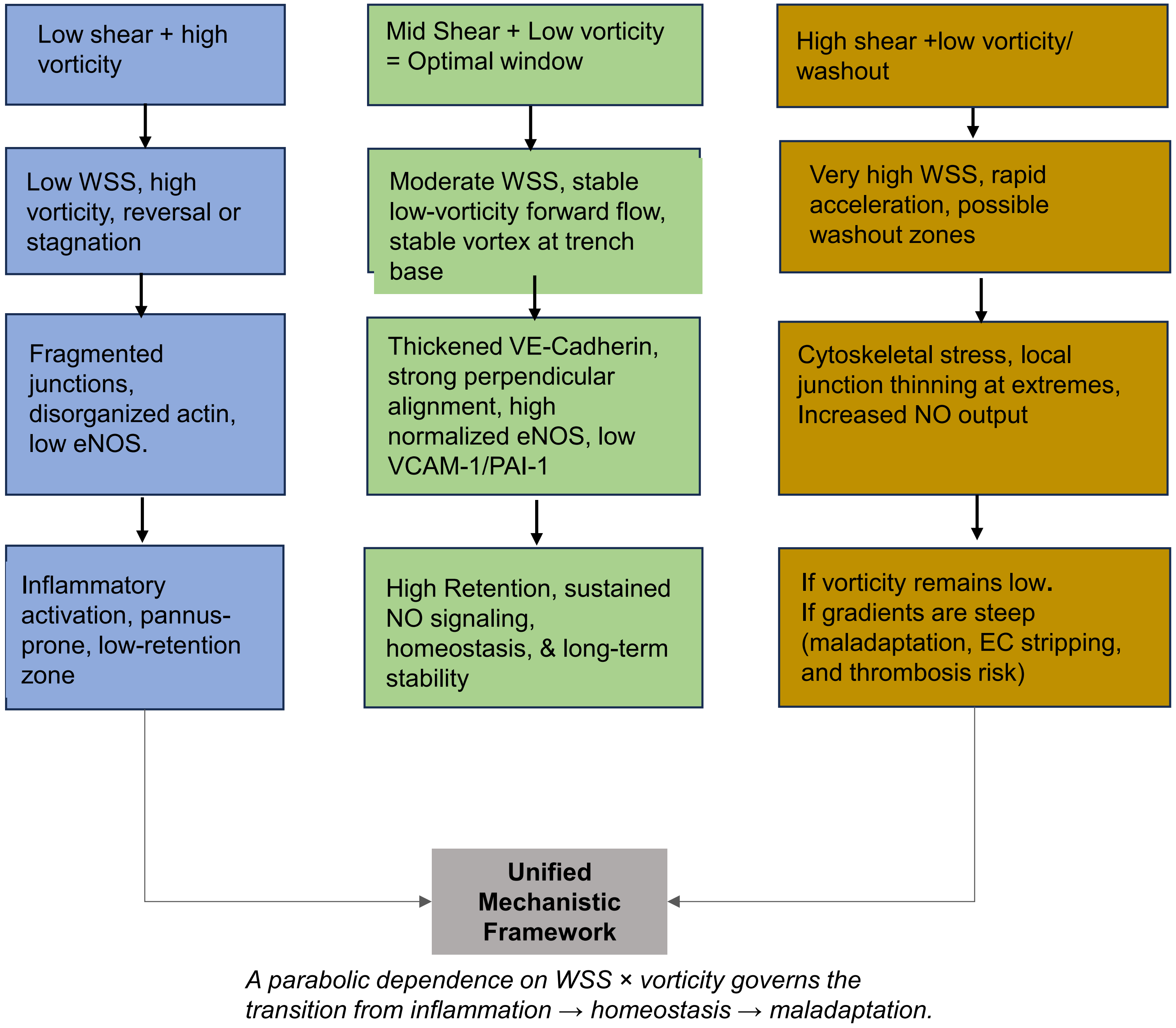


**Supplementary Fig. 10 | Unified mechanistic framework describing endothelial adaptation across shear–vorticity regimes.** Flowchart summarizing endothelial state transitions across shear–vorticity regimes. Low-shear/high-vorticity promotes inflammatory remodeling; mid-shear/low-vorticity supports homeostasis; high-shear/low-vorticity washout induces maladaptation when gradients are steep. A parabolic dependence on WSS × vorticity governs these transitions.

| Marker | Model | $R^2$ | Adjusted $R^2$ | Notes |
|---|---|---|---|---|
| **Coverage** | Linear | 0.55 | 0.51 | |
| | **Quadratic** | 0.62 | 0.57 | **Best fit** |
| | Logistic | --- | --- | Bounded |
| **Coherency** | Linear | 0.25 | 0.17 | |
| | Quadratic | 0.31 | 0.21 | **Best fit** |
| **Thickness** | Linear | 0.14 | 0.05 | |
| | Quadratic | 0.30 | 0.20 | **Best fit** |
| **Alignment** | Linear | 0.8 | 0.78 | |
| | **Quadratic** | 0.85 | 0.83 | **Best fit** |
| **eNOS** | Linear | 0.58 | 0.54 | |
| | **Quadratic** | 0.61 | 0.56 | **Best fit** |
| **VCAM-1** | Linear | 0.09 | 0.00 | |
| | **Quadratic** | 0.16 | 0.05 | **Best fit** |
| **PAI-1** | **Linear** | 0.3 | 0.23 | **Best fit** |
| | Quadratic | 0.31 | 0.21 | |

**Supplementary Table 1 | Summary of model performance across biological outputs.** Comparison of linear, quadratic, and bounded logistic regression models used to relate endothelial responses to shear–vorticity coupling (WSS × vorticity). Endothelial coverage and coherency were logit-transformed prior to regression due to bounded behavior, while junction thickness, alignment, eNOS, VCAM-1, and PAI-1 were modeled in raw units. Quadratic models generally outperformed linear fits across most readouts, with the strongest performance observed for coverage ($R^2$ = 0.62), alignment ($R^2$ = 0.85), and eNOS ($R^2$ = 0.61). Logistic models are reported for bounded outcomes but do not yield $R^2$ values. PAI-1 was best described by a linear model.

| Readout | β1(WSS×Vort) | β2 (WSS×Vort)² | Optimum Zone ($-\beta_1/2\beta_2$) |
|---|---|---|---|
| **EC coverage** | 1.04E-06 | -6.03E-13 | 0.862E+06 |
| **Coherency** | 2.76E-07 | -7.03E-14 | 1.96E+06 |
| **Junction Thickness** | 4.93E-07 | -1.48E-13 | 1.67E+06 |
| **Alignment** | -8.40E-06 | 2.25E-12 | 1.87E+06 |
| **eNOS/area** | 9.48E-08 | -1.64E-14 | 2.89E+06 |
| **VCAM-1** | 1.48E-06 | -3.58E-13 | 2.07E+06 |
| **PAI-1** | 1.67E-07 | 0 | #DIV/0! |

**Supplementary Table 2 | Regression coefficients and mechanical optima derived from quadratic fits.** $\beta_1$ and $\beta_2$ coefficients corresponding to linear and quadratic dependence on WSS × vorticity are shown for each biological readout. Mechanical optima were computed as $-\beta_1/(2\beta_2)$. Optima converged within 0.8–2.9 × $10^6$ dyn·cm$^{-2}$·s$^{-1}$ for all metrics except PAI-1, consistent with a shared endothelial mechanoadaptation window.

| s/no | Antibodies | Host | Dilution | Vendor | Catalog # |
|---|---|---|---|---|---|
| 1 | VE Cadherin (CD144) | Rat anti-Pig | 1:100 | Bio-Rad | MCA1748GA |
| 2 | eNOS | Rabbit anti-Pig | 1:100 | Cell Signaling | 9571L |
| 3 | VCAM-1 | Rat anti Mouse | 1:100 | Life Technologies | 14-1061-82 |
| 4 | PAI1 | Mouse anti Human | 1:100 | Life Technologies | MA140224 |
| 5 | Phalloidin-Alexa 635 | Chemical probe | 2.5uL/ml | Life Technologies | A34054 |
| 6 | CD31 (PECAM-1) | Mouse anti-pig | 1:100 | Life Technologies | MA5-18135 |
| 7 | Nuclei (DAPI) | - | 2 drops per mL | Life Technologies | R37606 |

**Supplementary Table 3 | Antibody List.** Details of all antibodies and chemical probes used to detect VE-cadherin, eNOS, VCAM-1, PAI-1, CD31, and F-actin. The table reports host species, working dilutions, vendors, and catalog identifiers. All antibodies were selected based on validated or reported cross-reactivity with porcine endothelial cells. Phalloidin is listed as a chemical probe without host species.

| Parameter | 0° trench | 22.5° trench | 45° trench |
|---|---|---|---|
| **Nodes** | 60,266 | 60,266 | 32,704 |
| **Elements** | 301,571 | 301,571 | 162,777 |
| **Element type** | Tet4 | Tet4 | Tet4 |
| **Global element size** | $3 \times 10^{-5}$ m | $3 \times 10^{-5}$ m | $3 \times 10^{-5}$ m |
| **Minimum edge length** | $1.05 \times 10^{-4}$ m | $1.05 \times 10^{-4}$ m | $4.8 \times 10^{-5}$ m |
| **Maximum aspect ratio** | 15.495 | 15.495 | 14.048 |
| **Minimum orthogonal quality** | 0.020 | 0.020 | 0.028 |
| **Inflation layers** | 5 | 5 | 5 |
| **Inflation growth rate** | 1.2 | 1.2 | 1.2 |

**Supplementary Table 4 | CFD mesh resolution and quality for all microtrench geometries.** All meshes were generated in ANSYS Meshing using quad-dominant surface meshing and swept tetrahedral elements (Tet4). Five inflation layers (growth rate 1.2) were applied to all trench surfaces. Mesh quality was assessed using standard ANSYS Meshing metrics; all meshes produced numerically steady solutions without nonphysical oscillations.

(a)

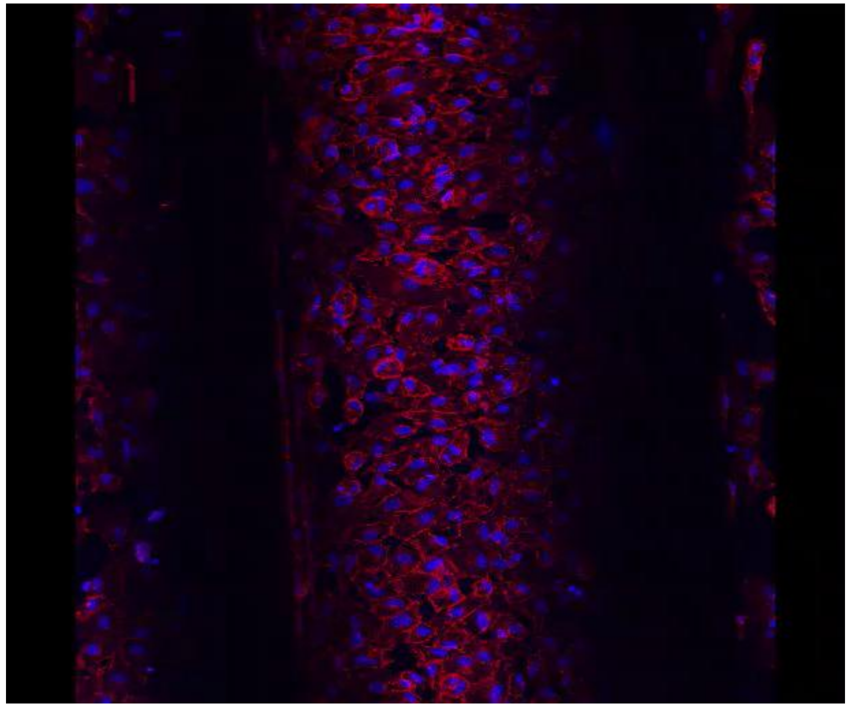

(b)

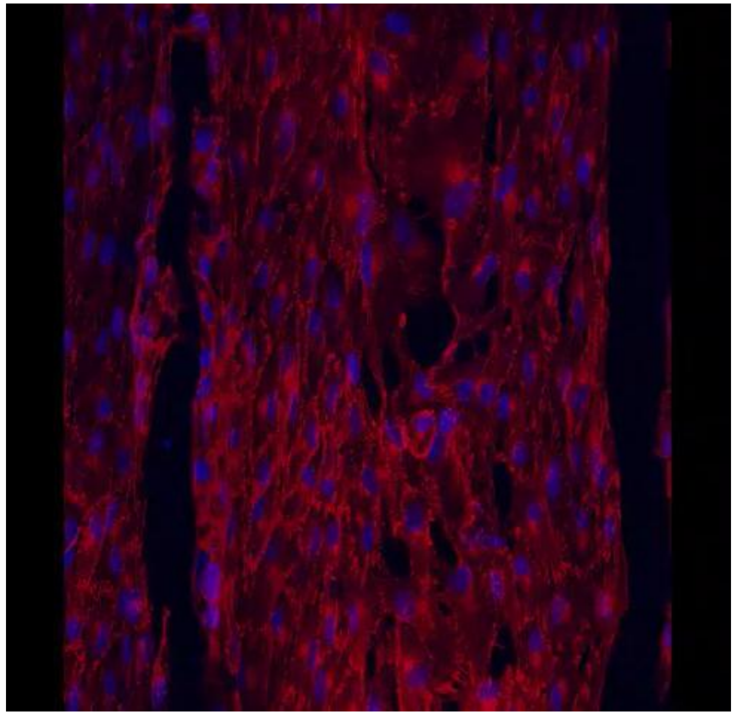

(c)

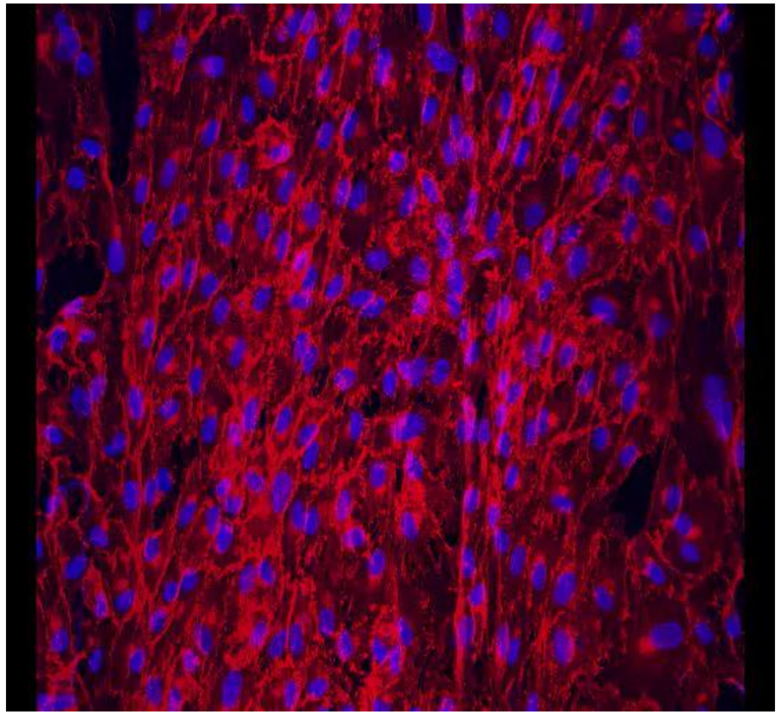

**Supplementary Movies | (a)** 0° trench showing endothelial retention after 48h of shear. (b) 22.5° trench showing endothelial retention after 48h of shear (c) 45° trench showing endothelial retention after 48h of shear
